# Interactions between circuit architecture and plasticity in a closed-loop system

**DOI:** 10.1101/2022.11.23.517718

**Authors:** Hannah L. Payne, Jennifer L. Raymond, Mark S. Goldman

## Abstract

Determining the sites of plasticity underlying changes in neural activity and behavior is critical for understanding mechanisms of learning. Identifying such sites from neural recording data can be challenging due to feedback pathways that impede reasoning about cause and effect. We studied interactions between feedback, neural activity, and plasticity in the context of a closed-loop motor learning task for which there is disagreement about the loci and directions of plasticity. We constructed a set of models that differed in the strength of their recurrent feedback. Despite these differences, each model successfully fit a large set of neural and behavioral data. However, the patterns of plasticity predicted by the models fundamentally differed, with the sign of plasticity at a key site changing from depression to potentiation as feedback strength increased. Guided by our analysis, we suggest how such models can be experimentally disambiguated. Our results address a long-standing debate regarding cerebellum-dependent motor learning and demonstrate how learning-related changes in neural activity can appear to contradict the sign of the underlying plasticity when feedback is present.

## 1 Introduction

Synaptic plasticity distributed across multiple sites of a circuit is thought to underlie changes in behavior. To understand how such plasticity supports learning, it is necessary to identify sites of plasticity, determine how this plasticity produces changes in neural activity, and link these changes to behavior. Thus far, such characterization has proven difficult in the vertebrate brain.

A key challenge is the ubiquity of feedback loops in neural systems. Feedback loops can be internal to the brain — as in the recurrent circuits thought to underlie short-term memory, predictive coding, and gain control (Constantinidis & Wang, 2004; Douglas & Martin, 2007; Vogels et al., 2005) — or partially external to the brain, arising whenever a behavior influences sensory input from the environment that, in turn, influences subsequent behavior, such as during control of movements (Robinson, 1965; Sperry, 1950; Todorov & Jordan, 2002; von Holst & Mittelstaedt, 1950). Such feedback loops make it challenging to identify the sites of plasticity that underlie learning. Because direct measurement of synaptic strength is extremely difficult in intact behaving animals, a common approach is to instead infer changes in synaptic strength from observed changes in neural firing (for example, Gilbert & Thach, 1977; Moita et al., 2003; Yao & Dan, 2001). As a simplifying assumption, neural systems are commonly treated as if they are feedforward circuits, with changes in neural activity at a particular site attributed to plasticity somewhere upstream of that site (**Figure 1A**). However, in systems with feedback loops, the distinction between upstream and downstream is ill-defined, so that changes in the activity of a neuron after learning do not necessarily reflect plasticity in its nominally upstream inputs (**Figure 1B**). Thus, feedback loops can confound the inference of changes in synaptic strength from observed changes in neural activity.

**Figure 1:**
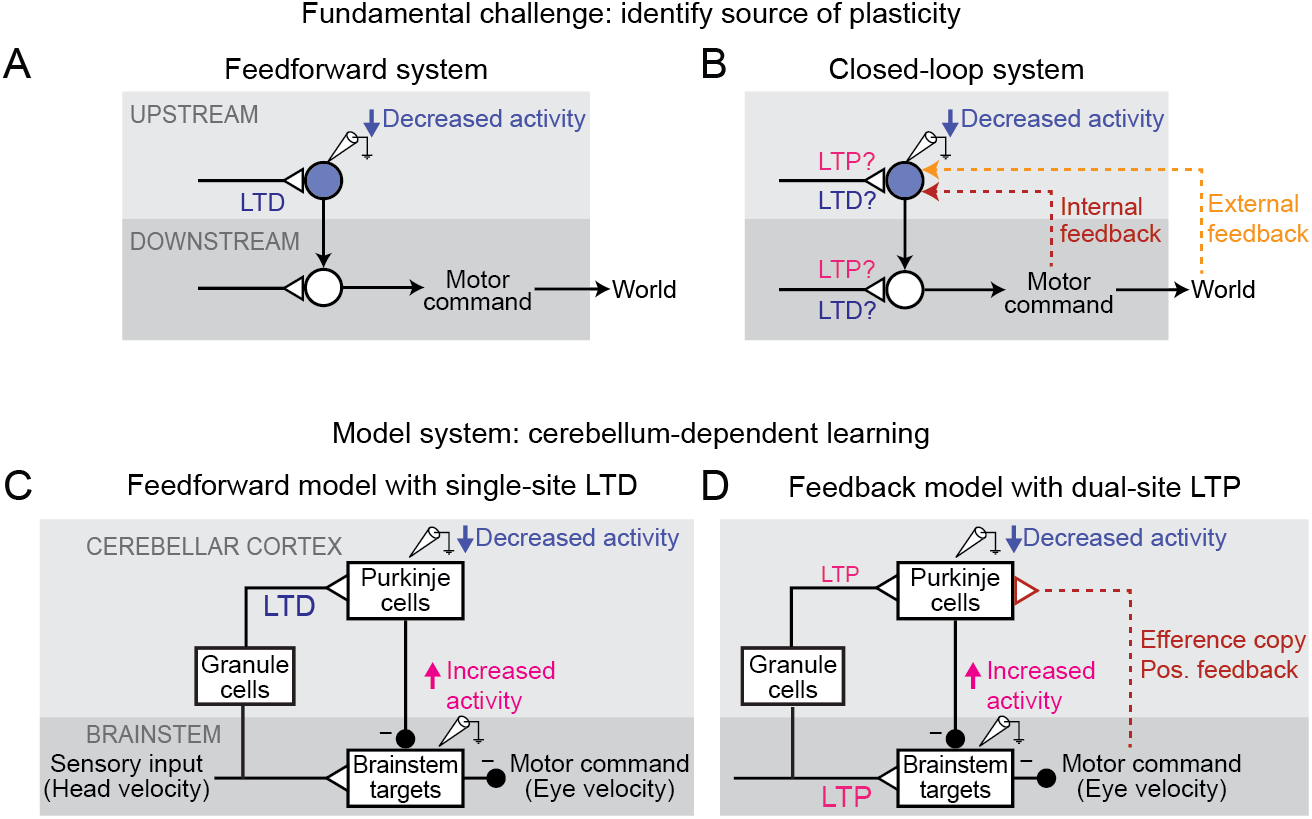
Feedback can obscure plasticity in neural systems. **(A)** Purely feedforward circuit. A decreased neural response to a stimulus after learning can be attributed to LTD of excitatory synapses (or equivalently, LTP of inhibitory synapses) upstream of the recorded neuron. *Triangles*, plastic synapses. **(B)** Recurrent circuit, with both internal and external feedback. The same decreased neural response can no longer be definitively attributed to plasticity at nominally upstream sites. Instead, plasticity at inputs to a second site that is a postsynaptic target of the recorded neuron, and thus appears to be downstream, may feed back to affect the recorded neuron’s activity. Such “downstream” plasticity may mask the effects of “upstream” plasticity. **(C)** Feedforward circuit hypothesized by Marr, Albus, and Ito to support motor learning. Before learning, vestibular (head velocity) inputs to the brainstem drive compensatory eye movements that are directed opposite to rotation of the head. After learning, LTD of vestibular inputs to Purkinje cells reduces inhibition onto brainstem neurons, increasing eye movement amplitude. **(D)** Simplified version of the feedback circuit hypothesized by Miles and Lisberger (1981) to underlie motor learning. After learning, LTP of vestibular inputs to the brainstem drives larger contraversive eye movements during the VOR in the dark. Efference copy of these eye velocity commands (*red pathway*) leads to decreased Purkinje cell activity, despite LTP of the vestibular input to Purkinje cells.

Such interactions between circuit feedback and plasticity are at the heart of a decades-long debate about how the cerebellum implements motor learning. The classic Marr-Albus-Ito model assumes a feedforward architecture in which errors are reduced through changes in the synaptic inputs to Purkinje cells, the sole output neurons of the cerebellar cortex (**Figure 1C**; Marr, 1969; Albus, 1971; Ito and Kano, 1982). This is consistent with a large number of studies suggesting that long-term depression (LTD) occurs at the excitatory parallel fiber inputs to Purkinje cells in response to error signals carried by climbing fiber inputs, effectively implementing negative reinforcement learning (“parallel fiber-Purkinje cell LTD”; Gilbert and Thach, 1977; Ito and Kano, 1982; Mauk, Steinmetz and Thompson, 1986; Sakurai, 1987; Coesmans *et al*., 2004; Medina and Lisberger, 2008; Kimpo *et al*., 2014; Inoshita and Hirano, 2018; Jang, Shim and Kim, 2020, but see Schonewille *et al*., 2011). In contrast, other experimental observations raised the possibility that the learning-related changes in Purkinje cell firing could instead be due to feedback of changes occurring *outside* of the cerebellar cortex (“Miles-Lisberger hypothesis,” **Figure 1D**; Miles and Lisberger, 1981). Furthermore, these experiments were interpreted as evidence that parallel fiber-to-Purkinje cell plasticity within the cerebellar cortex was in the opposite direction (long term potentiation, LTP) from the LTD predicted by the Marr-Albus-Ito model. These opposing conclusions about the sites and directions of plasticity underlying cerebellum-dependent motor learning remain unreconciled.

Here we use a data-driven, computational approach to determine how the strength of feedback in a circuit determines the sites and the direction of synaptic plasticity required during learning to accomplish a given change in behavior. We aggregated neural and behavioral data from a large set of experiments testing oculomotor performance and learning, and fit a series of computational models that systematically differed in the level of internal feedback. Our results provide a solution to a longstanding debate in the cerebellar field, and more broadly demonstrate how, in closed-loop systems, synaptic strength and neural activity at a given site can, counter-intuitively, change in opposite directions.

## 2 Results

### 2.1 Oculomotor learning circuit and closed-loop modeling strategy

We perform our studies within the context of the learned control of eye movements. Oculomotor learning provides a powerful experimental system because it regulates a relatively simple transformation from sensory inputs to motor output in a circuit whose anatomy and physiology have been extensively characterized. The oculomotor system generates eye movements to stabilize visual images on the retina during both motion of the body and motion of visual objects in the world. Vestibularly driven eye movements, known as the vestibulo-ocular reflex (VOR), stabilize gaze by counter-rotating the eyes in response to head movement and occur even in complete darkness. Visually driven eye movements, including “smooth pursuit” eye movements that track a moving target, stabilize images on the retina in response to visual input (Noda & Suzuki, 1979; Rambold et al., 2002).

Both the visual and vestibular functions of the oculomotor system are remarkably linear (Bagnall et al., 2008; Lisberger & Fuchs, 1978; McElvain et al., 2015; Payne et al., 2019; Walter & Khodakhah, 2006). Hence, we modeled this circuit with a network of linear temporal filters, *k*_YX_, each representing the transformation of a signal over a (mono- or multi-synaptic) anatomical neural pathway from node X to node Y (**Figure 2**). The net contribution of the excitatory and inhibitory synapses within each pathway is represented by a single linear filter that can have positive or negative value at any given time point. Vestibular sensory input encoding angular head velocity (H) drives eye movements (E) via a “direct pathway” through brainstem nuclei (*k*_EH_; **Figure 2**, *black*), and an indirect side loop through Purkinje cells (P) in the floccular complex of the cerebellar cortex (*k*_PH_; **Figure 2**, *black*) (Voogd et al., 2012). Purkinje cells also receive efference copy signals related to eye movement commands (*k*_PE_; **Figure 2**, *red*) and visual signals related to image motion (Voogd et al., 2012). The visual pathway is subdivided into a “retinal slip” (R) pathway, conveying, with a minimum 60 ms delay, motion of images across the retina (*k*_PR_, **Figure 2**, *orange*), and a “visual prediction” (T) pathway, providing non-delayed information about predictable target motion (*k*_PT_, **Figure 2**, *orange*). Signals conveying visual predictions have been recorded in cortical pathways that provide input to the floccular complex of the cerebellar cortex (Ilg & Thier, 2008) and, while not needed to explain the main qualitative results of the paper, were included to account for oculomotor tracking of predictable visual targets with no delay or in the absence of sustained retinal slip (**Figure 3–Figure Supplement 1**; Becker & Fuchs, 1985; Leung & Kettner, 1997; Stone & Lisberger, 1990) as in previous models of smooth pursuit (Barnes & Asselman, 1991; Orban de Xivry et al., 2013). Finally, neurons in the vestibular nucleus of the brainstem combine inhibitory input from the Purkinje cells (*k*_EP_, **Figure 2**, *black*) with direct vestibular input (*k*_EH_; **Figure 2**, *black*), and project to motor neurons in the abducens and oculomotor nuclei to control eye velocity.

**Figure 2:**
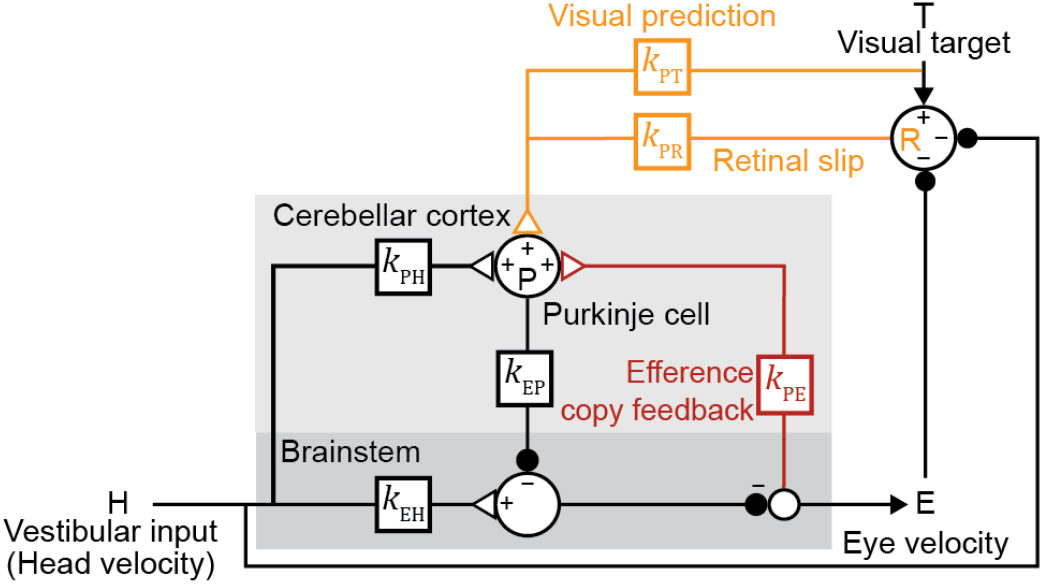
Linear filter circuit model. Signal transformations between different nodes of the circuit are modeled as linear filters *k*_XY_ (*boxes*). Purkinje cell population firing rate (P) is driven by vestibular stimulation (head velocity, H) through *k*_PH_, by efference copy of eye movement commands (E) through *k*_PE_ (*red*), and by visual signals through *k*_PR_ (*orange*, retinal slip velocity) and *k*_PT_ (*orange*, predicted visual target velocity). Eye velocity is driven by a direct pathway carrying head velocity input to the brainstem (*k*_EH_) combined with inhibition from Purkinje cells (*k*_EP_). The neural circuits between the brainstem and the eye muscles, which compensate for the dynamics of the eye plant, are implicitly included in the filters *k*_EP_ and *k*_EH_. For each fixed strength of *k*_PE_, all other filters were fit to the data.

Understanding how each of these pathways contributes to learning requires identifying the signal transformations occurring in each pathway. This is challenging because the vestibular, visual, and efference copy signals are tightly correlated due to feedforward and feedback interactions. Previous models have attempted to address this issue by assuming a particular strength of efference copy feedback, or have quantitatively fit simpler open-loop models to limited sets of data that may not fully eliminate the confounds stemming from strongly correlated predictor variables (Blazquez et al., 2003; Hirata & Highstein, 2001; Lisberger, 1994; Tabata et al., 2002). Here, we fit an extensive set of Purkinje cell and eye movement data recorded before and after learning, while systematically varying the strength of efference copy feedback to combat the effects of correlations between pathways and enable solutions that have not previously been considered (see **Materials and Methods**). A regularization penalty encouraged smoothness of linear filters before learning and discouraged large changes in filter weights after learning.

### 2.2 Before learning: degeneracy of model fits

Linear filter models with any level of efference copy input, ranging from a feedback gain of zero (“No Feedback”) to one (“Strong Feedback”), closely accounted for the dynamics of neural activity and behavior before learning (**Figure 3**). The data shown in **Figure 3** consist of Purkinje cell and eye velocity responses to 25 different oculomotor stimulus conditions, including vestibular input alone (VOR), visual input alone (smooth pursuit), and combinations of visual and vestibular input, delivered as sinusoids or steps of stimulus velocity. Responses during additional frequencies of the VOR in the dark before learning were also included from datasets that include learning (see section “After learning: circuit feedback affects inferences about plasticity” and **Materials and Methods**). For every stimulus condition, both the No Feedback and Strong Feedback models fit the data similarly (**Figure 3–Figure Supplement 2**). Models with negative internal feedback also fit similarly (not shown), whereas positive feedback gains greater than one were not considered since such feedback causes instability. The strength of efference copy feedback to Purkinje cells was thus unconstrained by a large set of oculomotor data before learning.

**Figure 3:**
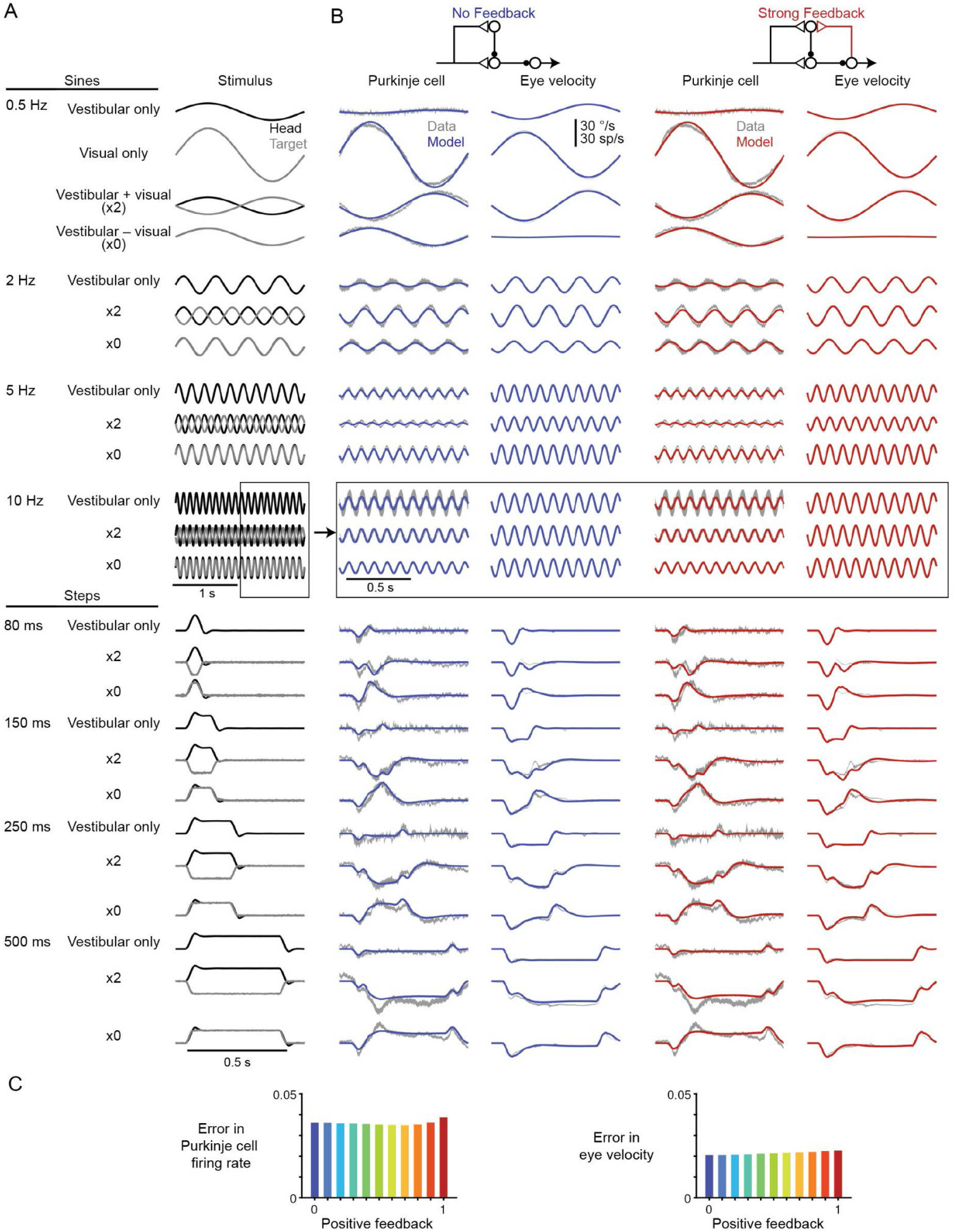
Models with or without efference copy feedback fit neural and behavioral data before learning. **(A)** Vestibular (“Head”, *black*) and visual (“Target”, *gray*) stimuli for each behavioral condition. Conditions consisted of vestibular input alone (“Vestibular only”; i.e. VOR in the dark), visual input alone (“Visual only”, i.e. smooth pursuit), vestibular input paired with oppositely directed visual input such that eye movements twice as large as normal were required to stabilize the image (“x2”), and vestibular input paired with visual input in the same direction such that eye movements must be eliminated to stabilize the image (“x0”, or VOR cancellation). Ipsiversive head and eye movements are plotted as positive values. **(B)** Purkinje cell firing rate and eye velocity measured experimentally (*gray*) and predicted by the No Feedback model (*blue*) or the Strong Feedback model (*red*). **(C)** Normalized root mean squared error of fits to Purkinje cell firing rate (*left*) and eye velocity (*right*) for all models.

To understand how models with vastly different efference copy feedback strengths could produce such similar outputs, we examined how other filters changed to compensate for the changing level of feedback. Some filters did not depend on feedback strength: the filters conveying vestibular input and Purkinje cell activity to the brainstem, *k*_EH_ and *k*_EP_, were nearly identical in all models, indicating that these pathways are well constrained by the data regardless of the strength of efference copy feedback (**Figure 4A**). In contrast, the filters carrying vestibular and visual inputs to Purkinje cells did vary with feedback strength. The vestibular input filter to Purkinje cells, *k*_PH_, changed from small and net negative in the No Feedback model to large and net positive in the Strong Feedback model (**Figure 4B**, *top*). Here, a positive filter weight indicates a net excitatory effect from an ipsiversive stimulus (e.g. increased excitation of a Purkinje cell in the right cerebellar hemisphere during head rotation to the right) and an inhibitory effect of a contraversive stimulus, whereas a negative filter weight indicates the opposite. The small net amplitude of the *k*_PH_ filter in the No Feedback model, evidenced by the small steady state step response (**Figure 4B**, *right*), directly reflects that Purkinje cells have minimal modulation of firing rate during the VOR. By contrast, in the Strong Feedback model, the same minimal modulation is achieved by a net positive *k*_PH_ filter that offsets the negative efference copy input through *k*_PE_. The retinal slip filter also varied with feedback strength, changing from acceleration-like to velocity-like as feedback strength decreased (**Figure 4B**, *bottom*) – this reflects that, in the Strong Feedback model, the efference copy feedback loop forms a temporal integrator that converts acceleration-like inputs to velocity-like outputs. The inferred strength and dynamics of the vestibular and visual inputs to the cerebellar cortex therefore are not well-constrained by this extensive set of data, and varied depending on the assumed strength of efference copy feedback, even before learning.

**Figure 4:**
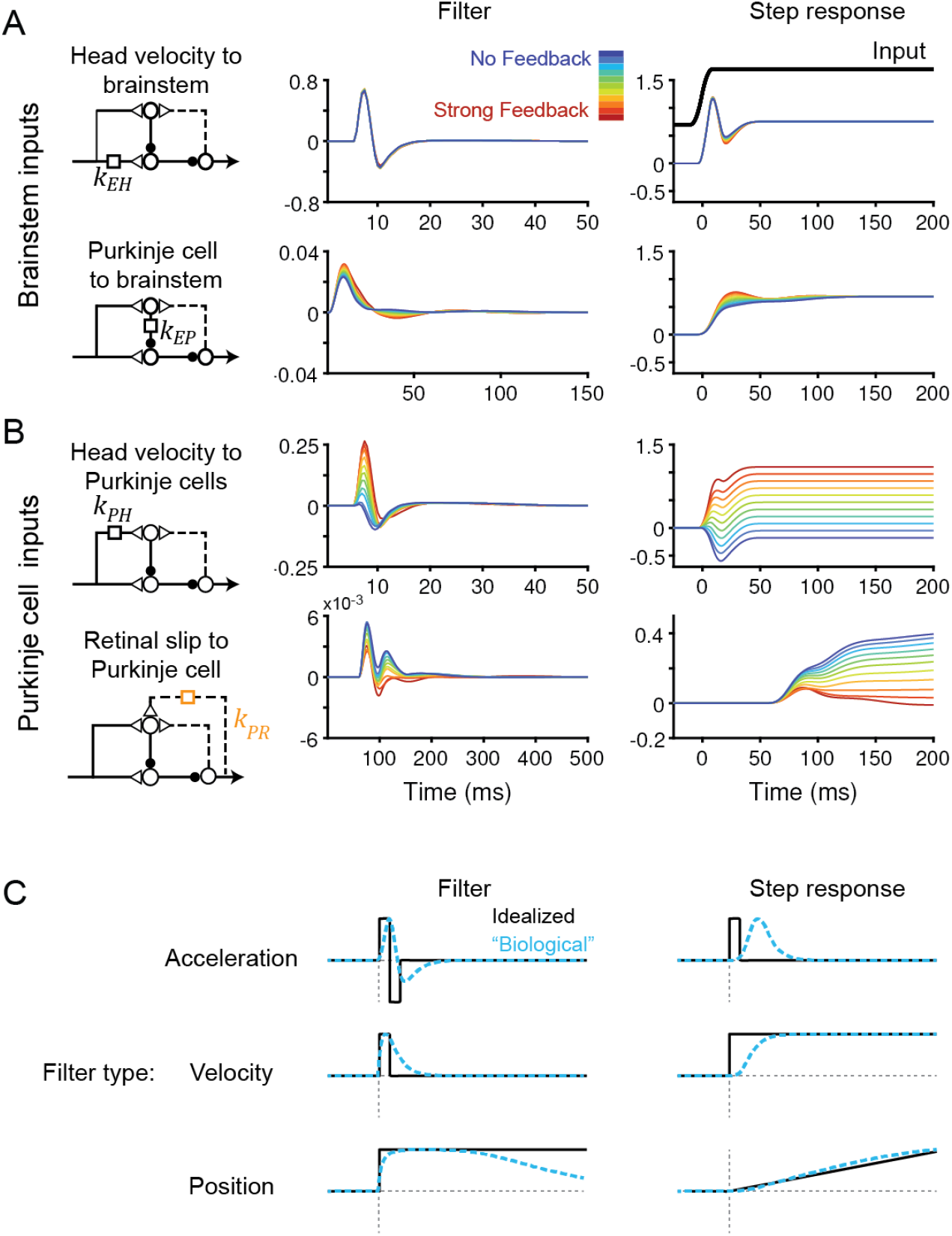
Temporal filters before learning are well-constrained for inputs to the brainstem, but not to Purkinje cells. **(A)** Filters conveying head velocity input (*top, k*_EH_) and Purkinje cell activity (*bottom, k*_EP_) to the brainstem for all models, ranging from No Feedback (*blue*) to Strong Feedback (*red*). The actual filter shape (*left*) and the response of the filter to a smoothed “step” input (*right*) are shown. Following Equation 1 (**Materials and Methods**), positive weights for *k*_EH_ cause oppositely directed (negative) changes in eye velocity. Most filters for *k*_EH_ are hidden beneath the trace for the No Feedback model. Units for step responses: °/s eye per °/s head (*top*), °/s eye per sp/s (*bottom*). For the linear filters, these units are multiplied by s^-1^. **(B)** Filters conveying head velocity (*top, k*_PH_) and retinal slip (*bottom, k*_PR_) input to Purkinje cells, as in (A). Units for step responses: sp/s per °/s head (*top*), sp/s per °/s retinal slip (*bottom*). **(C)** Schematic of idealized (*black*) and “biological” (*blue*) linear temporal filters and their corresponding step responses. Acceleration-like filters perform differentiation, velocity-like filters only change gain, and position-like filters perform leaky integration.

To understand the source of this apparent degeneracy, we analyzed a slightly simplified model with a single combined visual pathway that permits the analytic calculation of a closed-form solution for the model equations (**Materials and Methods**). This analysis revealed a strict degeneracy for some, but not all, parameters in the model. This is visualized in **Figure 5** for the steady-state component of the response by plotting the model cost function as the relevant filter parameters were varied. Consistent with the results of **Figure 4A**, the two brainstem pathways, *k*_EH_ and *k*_EP_, were fully constrained (**Figure 5A**), as illustrated by the single minimum in the cost function landscape. By contrast, there was a degenerate direction in parameter space for the three inputs to Purkinje cells: *k*_PH_, *k*_PE_, and *k*_PR_ (**Figure 5B)**, as illustrated by the flat valley in the cost function landscape, where the fit to the data was equally good for a range of different filter strengths. In this degenerate direction, *k*_PH_ and *k*_PE_ increase together while *k*_PR_ decreases. As discussed above, the concurrent increase in *k*_PH_ and *k*_PE_ indicates that the small Purkinje cell responses observed during the VOR in the dark could reflect either a small vestibular input (*k*_PH_) alone, or a large vestibular input offset by sufficient efference copy feedback (*k*_PE_). However, our analysis shows that the degeneracy is actually between all three (vestibular, efference copy, and visual) pathways rather than just the vestibular and efference copy feedback pathways. Ultimately, this degeneracy reflects that there are only two independently controllable variables, the vestibular and visual target stimuli, with eye velocity determined by these inputs. The practical implication of this analysis is that the vestibular, visual, and efference copy feedback filters to Purkinje cells cannot be fully determined using neural recording and behavioral data alone, but require an additional experimental strategy.

**Figure 5:**
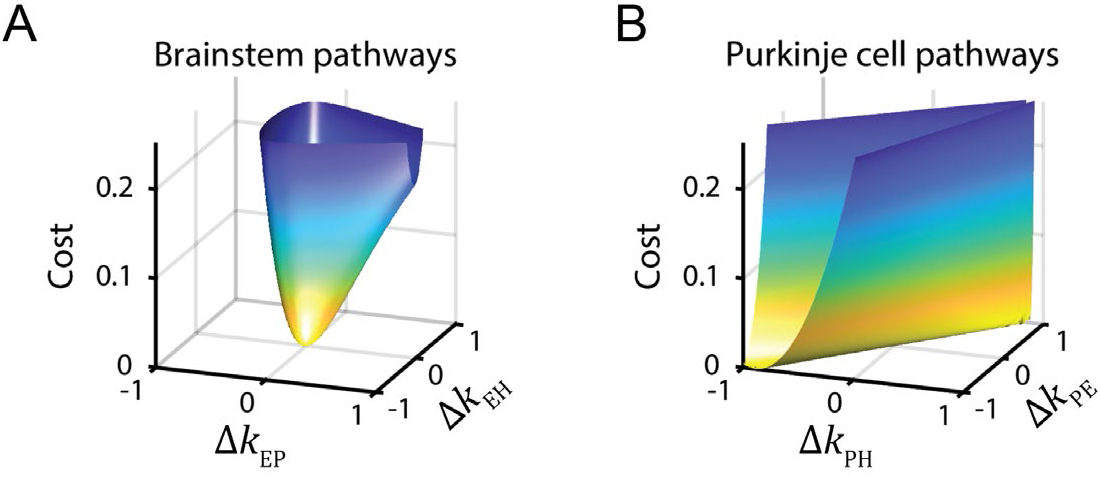
Cost function landscape illustrates degeneracy of input pathways to Purkinje cells, but not the brainstem. Degeneracy in the VOR circuit model fits, illustrated using a simplified model with merged retinal slip and visual prediction pathways (see **Materials and Methods**). Plots show the cost function “landscape” that is defined by the squared error between model output and experimental data. **(A)** Cost function landscape for the brainstem pathways (*k*_EP_ and *k*_EH_) has a single, well-defined minimum defining the best-fit parameter values. Δ*k*_XY_ indicates deviations from best fit values of filter responses at steady state (see **Materials and Methods**). **(B)** Cost function landscape for the Purkinje cell input pathways (*k*_PH_, *k*_PE_, and *k*_PR_) is degenerate, as reflected by the continuous valley of equally well-fit solutions. For each value of *k*_PH_ and *k*_PE_, the value of *k*_PR_ (not shown) was adjusted to minimize the cost.

### 2.3 After learning: circuit feedback affects inferences about plasticity

The fundamental problem of degeneracy before learning applies after learning as well, with critical implications for inferred learning mechanisms. Learning can increase or decrease the gain of the VOR. For simplicity, we focus below on learned increases in VOR gain unless otherwise specified; complementary changes occurred during learned decreases in VOR gain (**Figure 6–Figure Supplement 1 and 2**). We also, for ease of presentation, describe the response of the circuit to ipsiversive head turns (passive head rotation towards the side of the recorded neurons); the same arguments hold for contraversive head turns, but with opposite changes in firing throughout the VOR circuit.

A learned increase in VOR gain can be induced by pairing a vestibular stimulus with oppositely directed motion of a visual stimulus so that larger-than-normal eye movements are required to stabilize the image on the retina. Following learning, Purkinje cell responses to ipsiversive head turns in the dark *decrease* (Blazquez et al., 2003; Hirata & Highstein, 2001; Lisberger et al., 1994; Miles et al., 1980; Watanabe, 1985), which disinhibits brainstem target neurons, thereby increasing the amplitude of eye movements. Whereas these changes in behavior and neural activity are well-established, longstanding controversy concerns the sites and directions of plasticity underlying these changes.

The Marr-Albus-Ito model proposes that the observed decrease in Purkinje cell firing is caused by a decrease in the strength of vestibular input to Purkinje cells via LTD of vestibular parallel fiber-Purkinje cell synapses (**Figure 1C**) (Albus, 1971; Ito & Kano, 1982; Marr, 1969). However, the Marr-Albus-Ito model does not consider the potential contribution of efference copy input to Purkinje cells, and, in particular, the possibility that efference copy signals encoding the altered eye movements could contribute to the altered Purkinje cell firing after VOR learning. Later studies attempted to isolate the contribution of the vestibular input to Purkinje cell firing using a behavioral paradigm known as VOR cancellation, in which the eyes track a visual stimulus that moves exactly with the head, thus canceling the normal VOR eye movement response. During this paradigm, eye velocity in the orbit is close to zero – along with, presumably, any associated efference copy signals. Therefore, Purkinje cell activity during VOR cancellation was attributed to vestibular input alone. Surprisingly, Purkinje cell activity during VOR cancellation *increases* after learning – opposite to the *decrease* in activity (**Figure 1C,D**) observed during an identical vestibular stimulus presented in the dark (Lisberger, 1994; Miles & Lisberger, 1981). This increase in Purkinje cell activity during VOR cancellation was interpreted by Miles and Lisberger (1981) as evidence that the vestibular inputs to Purkinje cells must undergo potentiation during learning. The decrease in Purkinje cell activity observed during the VOR in the dark was then attributed to plasticity in the brainstem (*k*_EH_) that is relayed to the cerebellar cortex via efference copy feedback (*k*_PE_) (**Figure 1D**), rather than to LTD of vestibular parallel fiber inputs to Purkinje cells (*k*_PH_). However, a limitation of this hypothesis is its implicit assumption that visual input to the Purkinje cells was negligible during VOR cancellation because retinal slip velocity was close to zero. For an animal to ‘cancel the VOR’, it must use visual input to keep its eyes on the target; therefore, even retinal slip that is small but nonzero must have an influence somewhere in the oculomotor circuitry and cannot be discounted.

To assess these seemingly contradictory hypotheses, we examined the linear filters in our models before and after learning (**Materials and Methods**). To simulate learning, each model was fit to learned changes in behavior during the VOR in the dark across a broad range of stimulus frequencies from 0.5 Hz to 50 Hz (data from Ramachandran and Lisberger, 2005; **Figure 6A**) as well as to learned changes in Purkinje cell activity during the VOR at low frequencies (data from Lisberger et al., 1994; Watanabe, 1985). We compared the learning-related changes in filter shapes across models to assess how the strength of feedback in the circuit influences the inferred sites and directions of plasticity.

**Figure 6:**
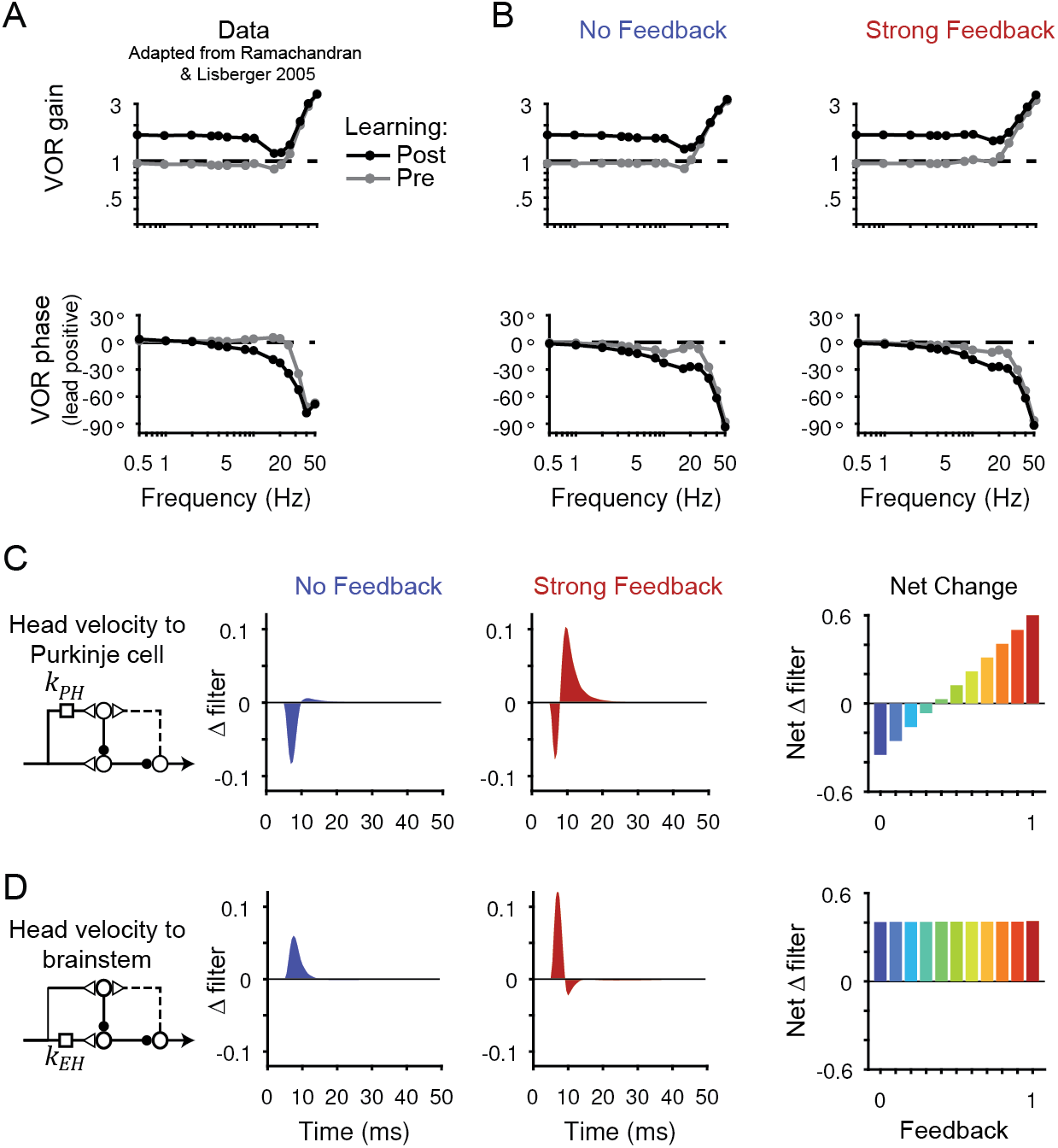
Circuit changes underlying learned changes in behavior. **(A**,**B)** Monkey (A) and model (B) eye velocity responses to sinusoidal vestibular input before (*gray*) and after (*black*) VOR learning. Behavior is quantified as the gain and phase of the eye relative to the head (0° phase represents the eye moving exactly opposite to the head). Data are adapted from Figure 4 of Ramachandran and Lisberger (2005). **(C)** *Left*, Dynamic change in the filter carrying head velocity input to Purkinje cells (*k*_PH_) after learned increases in VOR gain, for the No Feedback (*blue*) and Strong Feedback (*red*) models. Traces represent the *change* in filter strength after learning, rather than the absolute filter shape. *Right*, Net change in the filter *k*_PH_, calculated by numerically integrating the change in filter shape, for all models. Negative values indicate net depression and positive values indicate net potentiation. **(D)** Same as in (C) but for the filter carrying head velocity input to the brainstem (*k*_EH_). Note that the differences in filter shape between the No Feedback and Strong Feedback models in (D) largely reflect high temporal frequencies that were not well constrained by the experiments. Changes in filter shapes for intermediate feedback values and for learned decreases in VOR gain are shown in **Figure 6-Figure Supplement 2**.

All models, regardless of the strength of efference copy feedback, successfully reproduced VOR learning at the behavioral level (**Figure 6A,B, Figure 6-Figure Supplement 1)**. Each model captured the learned changes in gain and phase of the VOR observed in monkeys across stimulus frequencies (Ramachandran and Lisberger, 2005; **Figure 6A,B**). Despite these nearly identical changes in motor output, the underlying circuit plasticity depended critically on the strength of efference copy feedback. Similar to the baseline filters before learning, the net *changes* in the vestibular filters after learning depended on the level of feedback only for pathways in the cerebellar cortex (*k*_PH_, **Figure 6C**) and not in the brainstem (*k*_EH_; **Figure 6D**). Most strikingly, the No Feedback model displayed net depression of the vestibular input to Purkinje cells (*k*_PH_), whereas the Strong Feedback model displayed net potentiation at this site (**Figure 6C**, *right*) as well as depression of an acceleration-like component (Lisberger and Sejnowski, 1992; **Figure 6C**, *center*). Further, there was a gradual transition between these two extremes, with net depression switching to net potentiation at a feedback gain of around 0.4 (**Figure 6C**). Here, “depression” and “potentiation” refer to any combination of changes in excitatory and/or inhibitory inputs that result in decreases and increases, respectively, in the postsynaptic response and do not attempt to disambiguate, for example, between LTD of excitatory inputs and LTP of inhibitory inputs onto the same site (Carey, 2011). We confirmed that the direction of net plasticity of *k*_PH_ in each model was necessary by showing that the models failed to reproduce the experimentally observed changes in neural activity during the VOR after learning when the corresponding direction of plasticity was blocked (**Figure 6-Figure Supplement 3**).

The switch from net depression to net potentiation at *k*_PH_ with increasing feedback strength arises naturally from the model equations. For a given level of positive feedback strength *k*_PE_ (see **Discussion** for predicted effects of allowing *k*_PE_ to change over learning), the learning-related changes in Purkinje cell firing during the VOR in the dark gives:

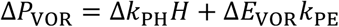

For models without feedback (*k*_PE_ = 0), depression of *k*_PH_ is required to reproduce the experimentally observed decrease in Purkinje cell firing during the VOR in the dark (Δ*P*_*VOR*_ < 0 requires Δ*k*_*PH*_ < 0; **Figure 6–Figure Supplement 4A-C**). For models with feedback (*k*_PE_ > 0), the learned change in eye velocity Δ*E*_VOR_ will contribute to Δ*P*_VOR_. The sign of this contribution is negative (since the eyes rotate even faster opposite to the head after learning) and increases in size with increasing feedback strength *k*_PE_. For sufficiently large feedback strength *k*_PE_, the term Δ*E*_VOR_*k*_PE_ will be more negative than Δ*P*_VOR_; hence, the change in the vestibular filter Δ*k*_PH_ must be positive to produce the observed change in Purkinje cell firing (**Figure 6–Figure Supplement 4D-E**). Thus, both the No Feedback and Strong Feedback models correctly reproduce the observed decrease in Purkinje cell activity, but they use oppositely directed changes in the vestibular pathway to do so.

### 2.4 Counterintuitive changes in neural activity after learning during VOR cancellation are consistent with multiple models

The observation that Purkinje cell activity during VOR cancellation increases after learning has previously been interpreted as evidence that the vestibular inputs to Purkinje cells (*k*_PH_) must undergo potentiation, rather than depression (Lisberger, 1994; Miles & Lisberger, 1981). To address whether such potentiation is indeed necessary, we examined the response during VOR cancellation of each model after learning. Note that the models were fit to reproduce learned changes in Purkinje cell activity only during the VOR in the dark, leaving changes in firing during VOR cancellation as a prediction of the model. In contrast to the previous interpretation, we found that all models – both those with potentiation and those with depression of kPH – successfully replicated this increase in Purkinje cell activity (**Figure 7**).

**Figure 7:**
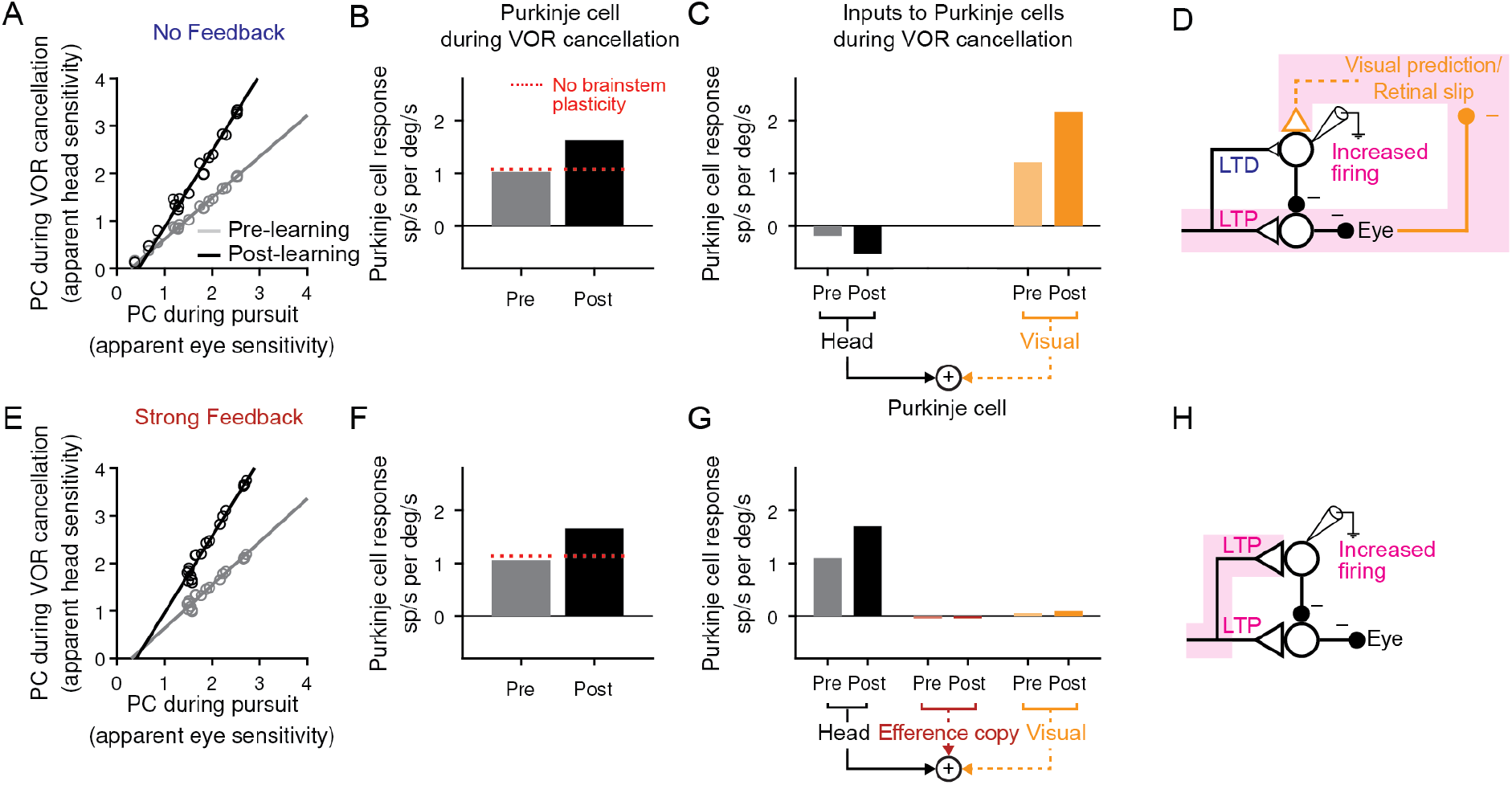
Explanation for changes in neural activity during VOR cancellation after learning. **(A)** Response of a population of simulated Purkinje cells during VOR cancellation compared to response during visual pursuit before (Pre) and after (Post) VOR learning for the No Feedback model. Values shown are in units of sp/s per °/s stimulus amplitude. Compare to Figure 13 of Lisberger *et al*. (1994). **(B)** Average response of Purkinje cell population in (A) during VOR cancellation. *Red dashed lines*, model results with brainstem plasticity blocked. **(C)** Inputs to Purkinje cells during VOR cancellation. The visual contribution represents the sum of both the retinal slip and visual prediction pathways. For the No Feedback model, the increase in input through the visual pathway after learning overshadows the decrease in input due to depression in the head velocity pathway. **(D)** Schematic illustrating that in the No Feedback model, the observed increase in Purkinje cell activity is due to negative feedback from vision. **(E,F,G)** Same as (A,B,C) for the Strong Feedback model. **(H)** In the Strong Feedback model, the observed increase in Purkinje cell activity is due to potentiation of vestibular inputs to Purkinje cells. Visual and efference copy pathways not shown because their contributions to Purkinje cell activity was minimal.

In the Strong Feedback model, the learned increase in Purkinje cell firing rate during VOR cancellation is not surprising: the vestibular input to Purkinje cells (*k*_PH_) potentiates, directly driving the increase in firing rate (**Figure 7E-H**). Because eye velocity was small during VOR cancellation and was strongly encouraged to not change during learning (Guo and Raymond, 2010; **Materials and Methods**), efference copy input to Purkinje cells made essentially no contribution to the increase in Purkinje cell activity after learning.

More surprisingly, the No Feedback model could also reproduce the observed increase in Purkinje cell activity during VOR cancellation after learning (**Figure 7A,B)**, despite net *depression* of the vestibular input to Purkinje cells (*k*_PH_) (**Figure 6, Figure 7C**). This was accomplished by the second feedback pathway in the circuit: external negative feedback of visual error, which drives corrective motor commands (**Figure 7C,D**). This visual feedback loop allows the VOR to be nearly completely canceled, despite plasticity in the brainstem (*k*_EH_) and cerebellar cortex (*k*_PH_) that would otherwise drive larger-than-normal eye movement commands in response to vestibular input. In essence, the visual feedback loop “works harder” after learning to counteract the increased VOR gain. It does this by increasing the activity of Purkinje cells, which then inhibit their target neurons in the brainstem to suppress the VOR. The increase in activity was predominantly due to the visual prediction pathway (**Figure 7C,D**) anticipating the regular sinusoidal pattern of eye movements and adjusting its output accordingly. However, while inclusion of the visual prediction pathways provided a closer quantitative match to the data, the core finding of increased Purkinje cell activity during cancellation did not depend upon visual prediction. In separate simulations that did not include a visual prediction pathway, Purkinje cell activity after learning increased during cancellation due to strong negative feedback of mildly increased retinal slip (**Figure 7–Figure Supplement 1**; note that *k*_PR_ was fixed in these simulations so the observed changes were not due to plasticity; see **Discussion** for predicted changes in *k*_PR_ with learning). Altogether, this analysis demonstrates that visual feedback can cause Purkinje cell activity during VOR cancellation to change in the opposite direction from the plasticity in the Purkinje cell’s vestibular inputs.

In both the No Feedback and Strong Feedback models, the correct learned increases in Purkinje cell activity during VOR cancellation depended on the existence of plasticity in the direct pathway through the brainstem. When the vestibular input to the brainstem (*k*_EH_) was held fixed during learning, motor learning was still successful at the level of the eye movement output during the VOR in the dark, but the learned increase in Purkinje cell firing during VOR cancellation no longer occurred (**Figure 7B,F**, *dashed lines*). This is because potentiation in the brainstem pathway (*k*_EH_) contributes to driving larger-than-normal eye movements in response to vestibular inputs and, during VOR cancellation, an increase in Purkinje cell activity is required to offset this. Therefore, in each model, two sites of plasticity – in the brainstem and in the cerebellar cortex – were required to explain the apparent paradox that learning decreases Purkinje cell firing measured during the VOR in the dark but increases firing measured during VOR cancellation.

### 2.5 Transient perturbation of activity distinguishes weak and strong feedback

The above results demonstrate that circuit feedback and circuit plasticity are interdependent – knowledge of one constrains the other. One approach that has been used to functionally assess circuit feedback is direct perturbation of neural activity (e.g., in models: Tsodyks *et al*., 1997; Sadeh and Clopath, 2020; in experiment: Pulizzi *et al*., 2016; Chettih and Harvey, 2019). A purely feedforward system should only respond briefly to transient stimulation, with a time course dominated by the intrinsic time constants of its neurons and synapses. In contrast, a system dominated by strong positive feedback should instead integrate transient stimulation, resulting in a prolonged response (Robinson, 1989). Mathematically, the response to such a perturbation of activity represents an independent constraint in addition to those provided by sensory-driven changes in activity. As a result, sensitivity analysis demonstrates that the previously observed degeneracy in the strengths of Purkinje cell inputs (**Figure 5C**) is eliminated when a “Purkinje cell stimulation” condition is added (**Figure 8A**). To leverage this constraint and quantitatively probe the strength of circuit feedback, model simulations were compared to the results of a previous study in which eye movements were evoked by electrical stimulation of floccular Purkinje cells in monkeys (Lisberger, 1994). In this experiment, brief stimulus trains (5 pulses at 200 Hz) applied in the absence of visual or vestibular stimulation evoked smooth eye velocity responses that decayed rapidly after stimulus offset (**Figure 8B**). We mimicked this experiment in our models by increasing Purkinje cell firing rate for 25 ms in the absence of vestibular or visual input. Models with weak or intermediate feedback produced brief eye velocity responses, akin to the data, whereas models with strong feedback produced prolonged eye velocity responses, unlike the data (**Figure 8C**). Although the precise strength of the efference copy feedback is difficult to ascertain because the time constant of decay of responses changes only gradually until the gain of the feedback loop approaches one, this suggests that feedback in this circuit is relatively weak (**Figure 8D**; see **Discussion**).

**Figure 8:**
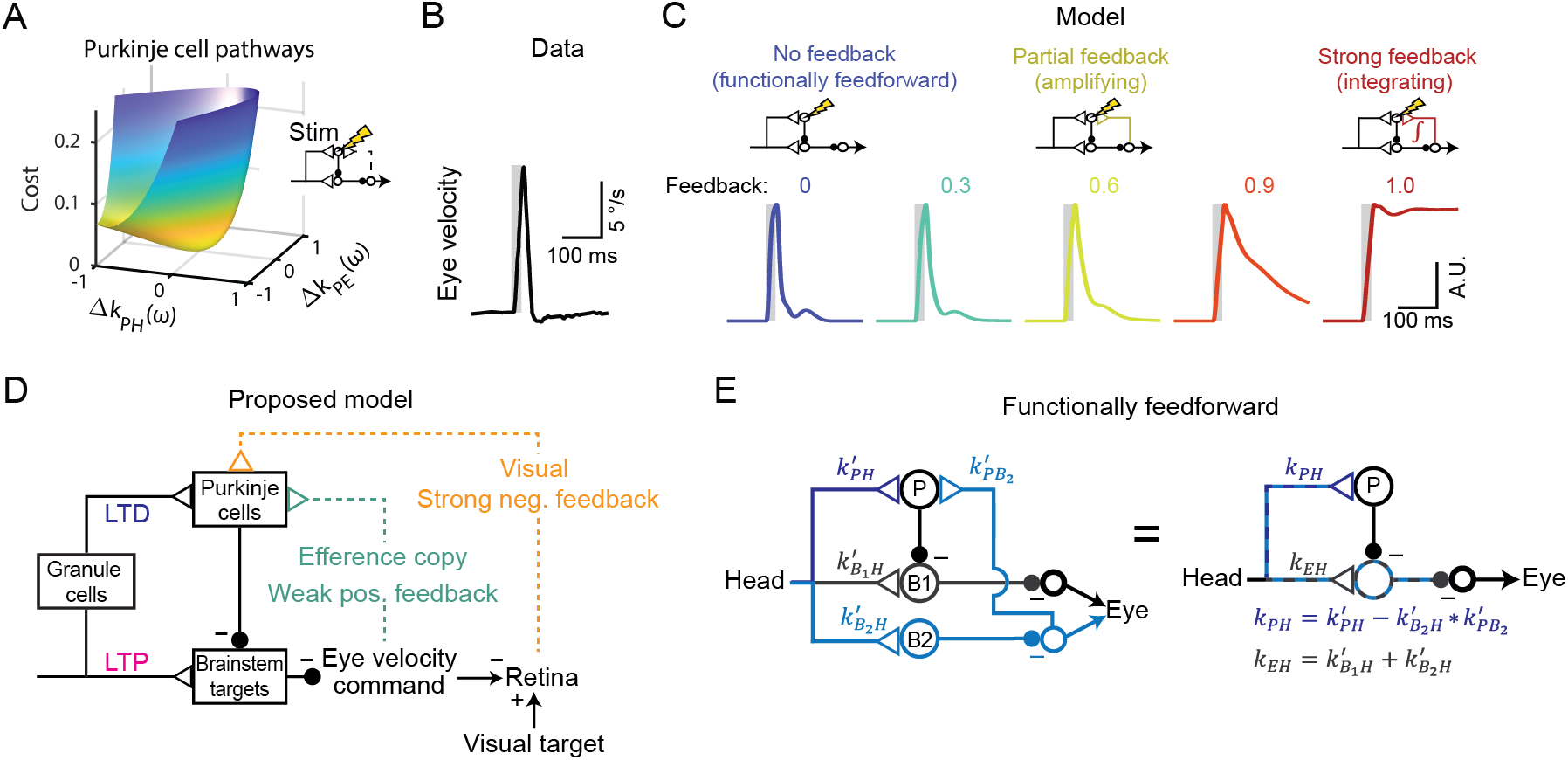
Circuit perturbations suggest functionally weak efference copy feedback. **(A)** Cost function landscape for the simplified model when a Purkinje cell stimulation condition is added. Compare to **Figure 5B. (B)** Empirically measured eye velocity in response to pulses of electrical stimulation in the Purkinje cell layer of the floccular complex (data replotted from Lisberger, 1994). **(C)** Model eye velocity responses to brief (25 ms) Purkinje cell stimulation, for models ranging from No Feedback (*left*) to Strong Feedback (*right*). **(D)** Proposed model reconciling previous experimental results. Efference copy feedback is weak to moderate, and both LTD in the cerebellar cortex and LTP in the brainstem pathway contribute to learned increases in VOR gain. In combination with brainstem plasticity, visual feedback drives the paradoxical increase in Purkinje cell activity during VOR cancellation after learning. (**E**) Feedforward architecture can exist despite anatomical feedback pathways between brain regions. *Left*, a circuit that is functionally feedforward despite anatomical projections that convey an efference copy from a population of brainstem neurons (B2) to the Purkinje cells (P). This circuit is purely feedforward, since the population of brainstem neurons that receive input from Purkinje cells (B1) do not loop back to the cerebellar cortex. *Right*, the equivalent feedforward circuit.

## 3 Discussion

Here we combine a systematic modeling approach with a wide array of neural and behavioral data to demonstrate how changes in neural activity and behavior arise from multiple sites of plasticity in a closed-loop circuit. Our results support the Marr-Albus-Ito model that LTD at the parallel fiber-Purkinje cell synapses contributes to VOR learning, because a range of models with such LTD and weak or no efference copy feedback could explain all recording and stimulation data. In addition, our results support the presence of a site of plasticity in the brainstem pathway, as proposed in the Miles-Lisberger model. This work reconciles these classic models by providing the first explanation of how the paradoxical increases in Purkinje cell activity during VOR cancellation can arise despite LTD of parallel fiber-Purkinje cell synapses. We thus provide a potential resolution of a longstanding debate about how the cerebellum implements motor learning, and exemplify more broadly how feedback can obscure underlying plasticity in even a relatively simple closed-loop system.

### 3.1 Net depression vs. potentiation of the parallel fiber-Purkinje cell synapses during learning

The most fundamental difference between models with different feedback strengths was the net direction of plasticity of vestibular inputs onto Purkinje cells: depression in weaker feedback models, and potentiation in stronger feedback models. Net depression at this site is consistent with the classic model of cerebellar learning whereby elevated climbing fiber activity encoding motor errors drives depression of recently active parallel fiber inputs to Purkinje cells, thereby decreasing the Purkinje cell response to the same stimulus on the next trial and reducing future errors (Ito, 1982; Raymond & Medina, 2018). By contrast, net potentiation at this site contradicts the classic mechanism of climbing fiber-driven synaptic depression. Furthermore, when the gain of the feedback loop is one (Strong Feedback model), any change in the strength of the vestibular input to the Purkinje cells (*k*_PH_) must be precisely offset by a change in the strength of the vestibular inputs to the brainstem (*k*_EH_) to avoid runaway activity (Lisberger & Sejnowski, 1992; **Figure 8–Figure Supplement 1**). Our study therefore reconciles results from the VOR paradigm of cerebellar motor learning with those from the eye blink and smooth pursuit paradigms, where climbing fiber activity has been less controversially associated with depression of active parallel fiber inputs (Lisberger, 2021; Raymond & Medina, 2018).

### 3.2 Previous arguments for an efference copy positive feedback loop

A major implication of these results for the oculomotor system is that a strong efference copy feedback loop to the cerebellar cortex is not required to account for experimental observations. This feedback loop was originally proposed to explain several observations (Lisberger, 1994; Miles & Lisberger, 1981): (1) correlation of Purkinje cell firing with gaze velocity, including nearly zero modulation of Purkinje cell firing during the VOR, (2) maintenance of smooth pursuit in the absence of retinal slip, including during brief disappearance of a pursuit target behind an occluder, (3) paradoxical changes in Purkinje cell firing after learning during VOR cancellation, and (4) anatomical input to the cerebellum from brain areas carrying eye movement-related signals. We demonstrate in the **Results** that the first three of these observations can be equally well explained by models with weak or even no efference copy feedback to the cerebellar cortex. With regards to the fourth observation, although the nucleus prepositus hypoglossi and the medial vestibular nucleus provide mossy fiber afferents to the cerebellar flocculus (Escudero et al., 1996a, 1996b), these pathways could involve ‘spiraling’ connections of the cerebellum with these regions (**Figure 8E**) rather than closed feedback loops onto the same cells. Further, even if true feedback loops are present, their functional feedback strength could be weak, or even negative (**Figure 6C** and **Figure 8B,C**).

Later arguments for efference copy feedback arose from multiple linear regression analyses in which Purkinje cell activity was fit to linear combinations of position, velocity, and/or acceleration of vestibular input, retinal slip, and eye movement (“efference copy input”) (Blazquez et al., 2003; Hirata & Highstein, 2001). These analyses found non-zero values for the coefficients representing eye velocity, which has been interpreted as evidence for efference copy feedback to Purkinje cells. Such studies used only a single frequency sinusoidal stimulus, making position and negative acceleration predictors strongly correlated, and delays and phase shifts difficult to distinguish. More fundamentally, these models are a special case of the more general linear filter models analyzed here, for which we have shown that there is an inherent non-identifiability of the inputs to Purkinje cells.

### 3.3 Relation to other sloppy models

In complex biological circuits, many different configurations of biological parameters can enable a circuit to accomplish the same task, resulting in degeneracy in the relation between model parameters and circuit output (“sloppiness in model fits”) (Fisher et al., 2013; Foster et al., 1993; Goldman et al., 2001; O’Leary & Marder, 2016; Prinz et al., 2004; Transtrum et al., 2015). Previous work on such “sloppy models” has focused primarily on sloppiness in cellular or network dynamics. Here, we show how the presence of multiple feedforward and feedback pathways converging on the same site can lead to sloppiness in inferring the sites and signs of plasticity, but how additional data from perturbations may constrain the most fundamental aspects of this sloppiness.

### 3.4 Predicted changes in the distribution of plasticity before and after systems consolidation of cerebellar learning

Studies of the consolidation of cerebellar learning suggest that memories form first in the cerebellar cortex before being transferred to the brainstem or cerebellar nuclei (Kassardjian *et al*., 2005; Shutoh *et al*., 2006; reviewed in De Zeeuw, Lisberger and Raymond, 2021). The model presented here represents a single time point of learning, with most of the post-learning experimental data coming from animals that were trained on previous days as well as the day of recording (**Materials and Methods**). Due to this previous training, some consolidation of learning would be expected, consistent with the brainstem plasticity that we inferred. However, a different distribution of plasticity between the cerebellar and brainstem sites would be expected at other times. Before consolidation, learning-related decreases in Purkinje cell firing during the VOR may be supported solely by plasticity in the cerebellar cortex (Ito & Kano, 1982; Jang et al., 2020). If this is the case, then our model predicts that “paradoxical” increases in Purkinje cell responses occurring during VOR cancellation (**Figure 7**) would not be observed at the earliest stages of learning before brainstem plasticity has occurred. Following consolidation, if learning becomes fully independent of the cerebellar cortex, then this has interesting implications for the final sign of plasticity of the vestibular input to Purkinje cells (*k*_PH_): for the cerebellar cortex to return to its original, pre-learning output, either (1) if there is no efference copy feedback, the vestibular pathway onto Purkinje cells needs to reset to its original strength, resulting in zero net plasticity, or (2) if there is efference copy feedback, the vestibular and efference copy pathways need to provide exactly canceling inputs during performance of the VOR. In the latter case, in the absence of plasticity in the efference copy pathway, net LTP in the vestibular pathway (*k*_PH_) would be required to offset changes in input from the efference copy pathway due to brainstem plasticity. This provides the possibility that the net sign of plasticity in vestibular inputs to Purkinje cells could be LTD at early stages of learning and LTP after complete consolidation of learning.

### 3.5 Model assumptions

#### 3.5.1 Pathways undergoing plasticity

Our model only included plasticity in the pathways carrying vestibular information (*k*_PH_ and *k*_EH_), which have been the focus of debate about VOR learning. However, it is reasonable to expect that the efference copy and visual inputs to Purkinje cells may be governed by the same correlational learning rule thought to control plasticity at the vestibular parallel fiber-Purkinje cell synapses, in which elevated climbing fiber activity drives depression of recently active parallel fiber inputs to Purkinje cells. Applying this rule to the efference copy pathway, the observed timing of climbing fiber activity relative to eye movements during VOR learning (Raymond & Lisberger, 1998) would drive changes in *k*_PE_ in the correct direction to support learning (potentiation for learned increases in VOR gain; depression for learned decreases in VOR gain). However, large changes in *k*_PE_ would lead to alterations of VOR dynamics, as well as gain, due to changing the strength of the efference copy feedback loop.

For the visual pathway, the correlational learning rule implies potentiation of *k*_PR_ during both learned increases and decreases in VOR gain, because visual error simultaneously drives climbing fiber and parallel fiber input in the same direction, so parallel fiber-climbing fiber correlations are independent of the sign of visual error. Interestingly, this may explain the increase in Purkinje cell firing during smooth pursuit that is observed after *both* directions of VOR learning (Blazquez et al., 2003; Lisberger et al., 1994; Miles et al., 1980).

#### 3.5.2 Visual input

We modeled visual input only to the cerebellar cortex, motivated by studies that show large decreases in the amplitude of smooth pursuit eye movements following lesions of the floccular complex (Belton & McCrea, 2000; Burde et al., 1975; Estanol et al., 1979; Rambold et al., 2002; Westheimer & Blair, 1974; Zee et al., 1981). There is some anatomical evidence for direct visual input to the brainstem as well (Balaban, 1983), and our model could be extended to implement this. This additional visual pathway would make the inputs to the brainstem site degenerate when fitting to neural and behavioral data alone, similar to the degeneracy demonstrated among the three pathways onto the Purkinje cells. Models fit to data collected after complete lesions of the floccular complex, or synaptic blockade of synaptic input to Purkinje cells, could provide an additional constraint to resolve this degeneracy.

### 3.6 Experiments to further constrain the strength of efference copy feedback

The results presented in **Figure 8** suggest that efference copy feedback is relatively weak, consistent with LTD at the vestibular parallel fiber-Purkinje cell synapse supporting learned increases in VOR gain, but this leaves open a range of possible efference copy feedback strengths. The functional impact of feedback to Purkinje cells could be more directly assessed by comparing the motor response to Purkinje cell stimulation in the presence versus absence of parallel fiber input. One study in mice did just that by blocking granule cell activity, which appeared to have little effect on the dynamics of the motor response to Purkinje cell stimulation, supportive of a weak feedback model in mice, although this was not quantified (Wada et al., 2014). Separately, if efference copy inputs contribute similarly to Purkinje cell firing during both smooth pursuit and VOR in the dark, as they do in the Strong Feedback model, then selective inactivation of those inputs should affect both behaviors.

## 4 Acknowledgements

We thank Jay Bhasin for comments on the manuscript, and Salomon Muller and Emre Aksay for helpful discussions.

## 5 Author contributions

**H.L.P**.: Conceptualization, Methodology, Software, Formal analysis, Investigation, Writing - original draft preparation, Writing-review & editing, Visualization

**J.L.R**.: Conceptualization, Methodology, Investigation, Data curation, Writing - review & editing, Supervision, Funding acquisition

**M.S.G**.: Conceptualization, Methodology, Investigation, Writing - original draft preparation, Writing - review & editing, Supervision, Funding acquisition

## 7 Data and materials availability

Data and code will be made publicly available upon publication.

## 8 Materials and Methods

### 8.1 Data set

Four datasets were used to capture neural and behavioral dynamics before and after learning. First, neural and behavioral data before learning (“Dataset 1”; shown in **Figure 3**) were obtained from two male rhesus monkeys (*Macaca mulatta)* trained to perform a visual fixation task. A subset of this dataset has been published previously (Kimpo et al., 2014; Raymond & Lisberger, 1998). Briefly, neural responses were recorded extracellularly from Purkinje cells in the floccular complex of the cerebellar cortex while the monkeys made horizontal eye movements in response to various combinations of visual and vestibular stimuli. Vestibular stimuli consisted of passive whole-body rotation in the horizontal plane. Visual stimuli consisted of a horizontally moving target subtending 0.5° of visual angle, which was accompanied by a larger black-and-white pattern subtending 20° to 30° of visual angle for all stimulus conditions except for smooth pursuit. Four combinations of visual and vestibular stimuli were delivered: head movements in the dark (“Vestibular only,” which elicits the VOR), visual target motion with the head stationary (“visual only”, which elicits smooth pursuit eye movements), visual target and head motion at the same speed but in opposite directions (**Figure 3**, “vestibular + visual”, also referred to as ×2 because accurate tracking requires compensatory eye movements that are twice as large as normal), and visual target and head motion at the same speed and in the same direction (**Figure 3**, “vestibular – visual”, also referred to as ×0 because accurate tracking requires *no* rotation of the eyes in their sockets, and also known as “VOR cancellation” since normal VOR eye movements are suppressed). These combinations were delivered as sine waves in stimulus velocity with frequencies of 0.5 Hz, 2 Hz, 5 Hz, and 10 Hz at ±10 °/s, or as steps in stimulus velocity with durations of 80 ms, 150 ms, 250 ms, and 500 ms at 15 °/s. Smooth pursuit data were only available for 0.5 Hz sine waves (delivered at ±31.4 °/s), resulting in a total of 25 distinct conditions. Eye position (angle of the eye relative to the head) was measured using the scleral search coil method.

Eye velocity and neural activity from Dataset 1 were further processed as follows. Eye velocity was calculated by differentiating eye position. Saccades were removed from eye velocity traces using an automatic threshold algorithm. To allow comparison across datasets and with previous studies, we analyzed horizontal gaze velocity Purkinje cells (HGVPs), which are the largest subpopulation of floccular Purkinje cells and are widely considered to be important for horizontal gaze control (Katoh et al., 2015; Lisberger et al., 1994). Purkinje cells were considered HGVPs if (1) during smooth pursuit, firing rate was modulated by at least ±10 sp/s and the phase difference between peak firing rate and peak ipsiversive eye velocity was less than 45°; (2) during VOR cancellation, firing rate was modulated by at least ±10 sp/s and the phase difference between peak firing rate and peak ipsiversive head velocity was less than 45°; and (3) firing rate modulation was greater during horizontal than during vertical smooth pursuit. Purkinje cell firing rates were calculated by convolving raw simple spikes times with a 10 ms standard deviation Gaussian filter. Baseline firing rates were removed by subtracting a moving average calculated over an 11 s window. Eye velocity and Purkinje cell firing rates were then averaged across stimulus cycles for each cell. Neurons were only recorded on one side of the brain, so to account for the corresponding population in the opposite hemisphere, Purkinje cell responses to ipsiversive stimulation were averaged together with the inverted response to contraversive stimulation. Finally, data were averaged across all cells to create mean Purkinje cell firing rate and mean eye velocity traces for each stimulus condition.

The remaining datasets were used to fit changes that occurred after learning. Dataset 2 was taken from (Ramachandran & Lisberger, 2005) and consisted of VOR *behavior only* in *Macaca fuscata* before and after learning. In that study, the VOR was tested at 15 different frequencies from 0.5 to 50 Hz in three monkeys, both before learning and after several days of training to increase or decrease the gain of the VOR. We averaged the reported gain and phase of the eye movement response across all three monkeys **(Figure 6A)**. The 12.5 Hz point was excluded as an outlier since it was reported to likely be influenced by a mechanical issue and it was not seen in the one monkey tested using a more reliable vestibular stimulation technique (Ramachandran & Lisberger, 2005).

Datasets 3 and 4 consisted of changes in Purkinje cell *activity* after learning from (Lisberger et al., 1994; Watanabe, 1985). Both of these studies characterized the amplitude of Purkinje cell activity during the VOR before and after learning in monkeys (*Macaca fuscata* and *Macaca mulatta*, respectively), using 0.3 Hz sine waves (Watanabe, 1985) or steps (Lisberger et al., 1994) in vestibular stimulus velocity. Both the steady-state changes (averaged 100-200 ms after step onset) and transient changes (peak firing rate averaged 0-50 ms after step onset) after learning reported by Lisberger et al. were included in the cost function. Learning-related changes in neural activity at higher sinusoidal frequencies have not been reported. To account for different amounts of behavioral learning (change in VOR gain) in different studies, we normalized the change in Purkinje cell modulation by the change in VOR performance (1.36 sp/s per °/s head velocity per fractional change in VOR gain for VOR-decrease learning and −0.75 sp/s per °/s for VOR-increase learning, Lisberger *et al*., 1994; 0.72 sp/s per °/s for VOR-decrease learning and −0.55 sp/s per °/s for VOR-increase learning, Watanabe, 1985). The results from the two studies were averaged to yield target changes of 1.04 sp/s per °/s head velocity per fractional change in VOR gain during VOR-decrease learning, and −0.65 sp/s per °/s during VOR-increase learning. A change in behavior of 60% (e.g., VOR gain increase from 1.0 to 1.6) was simulated, comparable to the changes achieved in these studies.

### 8.2 Model implementation

#### Model equations

The model architecture was constrained by anatomy, similar to previous models (Clopath et al., 2014; Dean & Porrill, 2008; Lisberger, 1994; Lisberger & Sejnowski, 1992; Miles & Lisberger, 1981; Tabata et al., 2002; Yamazaki & Nagao, 2012) (**Figure 2**). In the direct pathway, vestibular signals (*H*) from the semicircular canals travel through Scarpa’s ganglion to excite the ipsilateral vestibular nuclei, which directly inhibit motor neurons in the oculomotor nuclei to produce contraversive eye movements (*E*). The motor output from this direct pathway is modified by the indirect pathway through the cerebellum, in which the vestibular signals travel through the cerebellar cortex via granule cells and Purkinje cells, before reuniting in the vestibular nucleus. In addition to vestibular signals, Purkinje cells in the floccular complex of the cerebellar cortex also receive visual and eye velocity-related signals as described in the main text. Note that Purkinje cells have an unusually high baseline firing rate (∼50–100 Hz); in our model, this baseline is subtracted, and bidirectional modulation of firing rate is represented by values above and below zero.

Based on evidence for linearity in the VOR circuit and cerebellar Purkinje cells (Bagnall et al., 2008; Lisberger & Fuchs, 1978; McElvain et al., 2015; Payne et al., 2019; Walter & Khodakhah, 2006), the model architecture is described by the following linear firing rate equations:

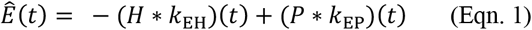

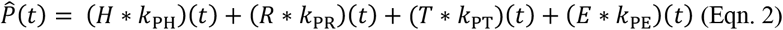

where * indicates temporal convolution, *E* is eye velocity, *H* is head velocity, *P* is Purkinje cell simple spike rate, *R* is retinal slip velocity, *T* is visual target velocity, and *k*_XY_ indicates the linear temporal filter to *X* from *Y*. *Ê*(*t*) and 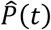 represent the model predictions for eye velocity and Purkinje cell firing rate.

The temporal filters *k*_EH_, *k*_EP_, *k*_PH_, and *k*_PR_ were parameterized using a raised cosine basis (Pillow et al., 2005) to efficiently capture biological filter shapes with relatively few parameters (10 basis vectors per filter for *k*_PH_, *k*_EH_, and *k*_EP_; 12 basis vectors for *k*_PR_). Each basis vector was given by:

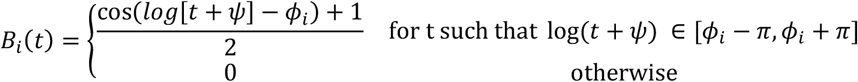

The time axis was logarithmically spaced so that the temporal filters were represented in greater detail at short timescales. The area of each basis vector was normalized to equal one. The duration of each filter was limited to *t*_max_ = 50 ms for *k*_PH_ and *k*_EH_, 150 ms for *k*_EP_, and 500 ms for *k*_PR_. These durations were chosen on the basis of preliminary model fits using longer durations, which nonetheless resulted in shorter optimal filters. The absolute minimum latency of each temporal filter was set to 5 ms for *k*_PH_ and *k*_EH_, and 1 ms for *k*_EP_, which resulted in similar latencies to the peak of the first basis vector of 7.0 ms for *k*_PH_ and *k*_EH_ and 7.5 ms for *k*_EP_, with latencies to the half-max of the first basis vector of 6.0 ms for *k*_PH_ and *k*_EH_ and 4.0 ms for *k*_EP_, generally consistent with reported latencies in the VOR circuit (du Lac et al., 1995; Lisberger, 1984). The absolute minimum latency of *k*_PR_ was set to 60 ms.

For the temporal filter conveying efference copy feedback to Purkinje cells, *k*_PE_, the temporal dynamics were represented by an exponential filter with a time constant of 3 ms, approximating a fast monosynaptic connection in order to allow the full range of frequencies of the VOR stimulus to be well-fit by the Strong Feedback model (**Figure 6A,B)**. The amplitude of this filter was set to make the total steady-state gain of the efference copy feedback loop vary from zero (“No Feedback”) to one (“Strong Feedback”) in steps of 0.1 to create a series of otherwise initially identical models. The total steady-state feedback gain corresponded to the time integral (or, in discretized time, sum) of the convolution of the two filters comprising the feedback loop, *k*_PE_ and 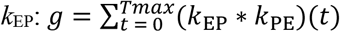. For each model, the relative strength of *k*_PE_ and *k*_EP_ could vary, but the total steady-state gain *g* was held fixed.

For the visual target pathway *k*_PT_, which represents visual prediction signals (sometimes referred to as “extraretinal” signals), we parameterized the filter as follows. For the experimental paradigms using sinusoidal visual stimuli, we fit the amplitude and phase of *k*_PT_ separately for each frequency in the data (0.5 Hz, 2 Hz, 5 Hz, and 10 Hz). For the paradigms using steps in visual stimulus velocity, *k*_PT_ was a single exponential filter with time constant 25 ms and a delay of 60 ms to account for the unpredictable nature of the step onset, and with amplitude as a free parameter that we fit. We note that the visual prediction filters for the sinusoidal and step stimuli are different from each other and were allowed to have different values before and after learning – this reflects our assumption that the visual prediction mechanism is able to nonlinearly adjust in order to account for different behavioral demands. The visual prediction pathway was included on the basis of several experimental observations. First, during smooth pursuit, primates can follow complex yet predictable visual stimuli with small (∼10 ms) or even negative (anticipatory) latencies (Leung & Kettner, 1997). Second, during smooth pursuit of a visual target that goes behind an occluder, eye velocity can change in anticipation of target reappearance, even anticipating expected changes in target velocity (Becker & Fuchs, 1985; Bennett & Barnes, 2004). Finally, anticipatory smooth eye movements can even occur in response to audio cues or stationary visual cues that predict upcoming target movement (Boman & Hotson, 1988; Jarrett & Barnes, 2005). Predictive visual signals have been observed at several sites that provide input to the floccular complex: in the medial superior temporal area (MST), a higher order region of motion processing in cortex (Kawano et al., 1994; Newsome et al., 1988; Sakata et al., 1983), in the dorso-lateral pontine nucleus (DLPN), a brainstem nucleus that projects to the flocculus (Mustari et al., 1988), and at mossy fiber inputs within the flocculus (Noda, 1986). In our model, if the visual prediction pathway is excluded, the main qualitative results (the directions and amplitudes of plasticity in the vestibular pathways over learning, and the explanation for the paradoxical changes in Purkinje cell activity) are unchanged, but the model fits during higher frequency visual stimulation become worse for all models, and the No Feedback model is unable to follow a target behind an occluder.

#### Model fitting

Model fitting was performed in two steps. First, the filters were initialized either using linear optimization with the model in open-loop configuration (i.e., with separate fits of Eqns. 1 and 2), or using the results of an adjacent model (i.e. in the Figures, the final fits for the model with g = 0 were used to fit the model with g = 0.1, the results from g = 0.1 were used to fit g = 0.2, etc.). Note that model fits were similar for both initializations, differing only in high frequency components that are not well-constrained by data. Second, the filters were fine-tuned using nonlinear optimization with the model in closed-loop configuration (i.e., with both Eqns. 1 and 2 operating simultaneously).

In the open-loop fitting step, we first performed two separate linear regressions corresponding to Eqns. 1 and 2 to provide initial estimates of the filters *k*_PH_, *k*_EH_, *k*_PR_, *k*_PT_, and *k*_EP_, using only data before learning (Dataset 1 and <= 10 Hz data before learning from Dataset 2). We then performed an additional pair of linear regressions corresponding to Eqns. 1 and 2 to provide initial estimates of the changes in the filters *k*_PH_ and *k*_EH_ after learning. To discourage overfitting, regularization was applied using Tikhonov matrices to minimize both the amplitude of the weights and the second derivative of the weights, with the following parameters. The regularization penalty for the first basis vector before learning equaled 1 for *k*_EH_, *k*_PH_, and *k*_PT_; 6 for *k*_PR_; 40 for *k*_EP_; and 0.25 for *k*_PT_. To encourage filters to be brief, the regularization penalties for *k*_PH_, *k*_EH_, and *k*_PR_ increased with the square of the basis index, so that longer timescale basis vectors were penalized more heavily than shorter timescale basis vectors. To keep the fit coefficients on a similar scale, each signal type used to fit the model (head velocity, target velocity, eye velocity, retinal slip velocity, and Purkinje cell firing rate) was normalized before fitting by dividing by its standard deviation, calculated across all 25 conditions before learning. Because the mean Purkinje cell firing rate was already subtracted, and the velocity signals were centered on zero, mean subtraction was not needed before normalization. The learned filters were converted back to real-world units after the fit was complete. To encourage filter weights after learning to be similar to those before learning, regularization was applied to the differences between the before-learning weights and the after-learning weights using the same Tikhonov matrices described above, but with all regularization penalties scaled ×12 to encourage relatively small changes in filter coefficients compared to the baseline values of the filter coefficients.

In the second, closed-loop fitting step, the filter coefficients were initialized to the values attained in the open loop step described above (or from the final results of an adjacent model) and then fine-tuned through a nonlinear fitting procedure conducted with the model in closed-loop configuration. The vestibular filters were fine-tuned separately from the visual filters to reduce the number of parameters that needed to be optimized simultaneously. First, the filters *k*_PH_, *k*_EH_, and *k*_EP_ before learning and the change in the filters Δ*k*_PH_ and Δ*k*_EH_ after learning were simultaneously fine-tuned in a single step using the MATLAB function *fmincon*. These filters were optimized according to the following cost functions:

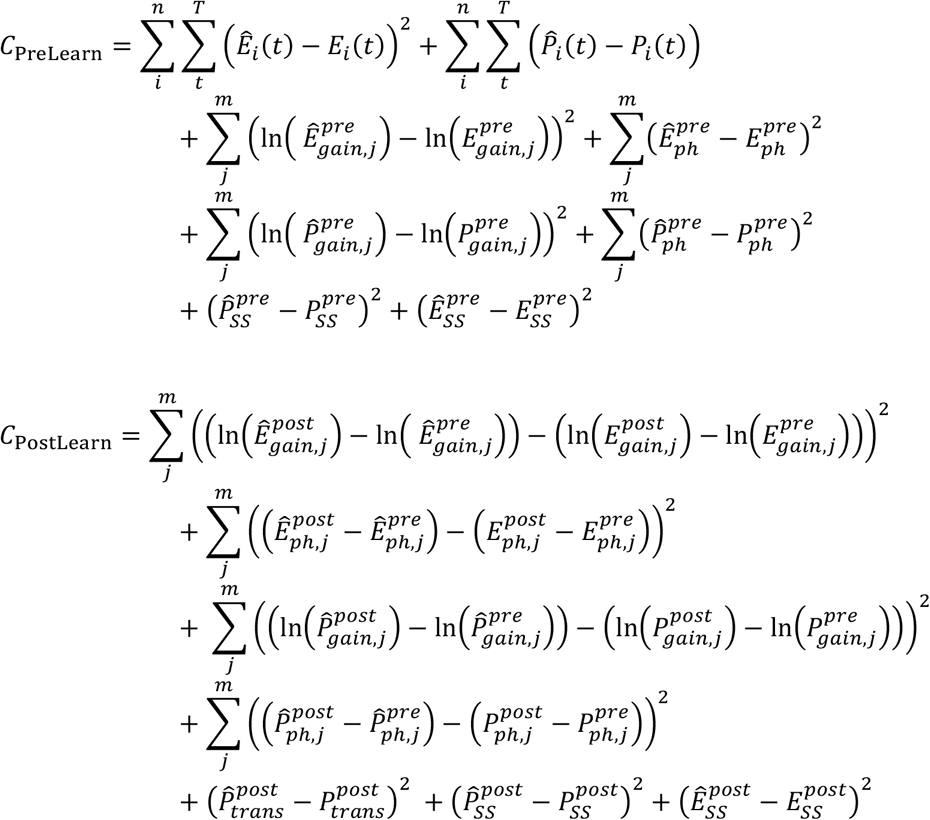

In *C*_PreLearn_, the first two terms correspond to fitting estimates (denoted by ^) to the eye velocity (*E*_i_) and Purkinje cell activity (*P*_i_) before learning for each of the *n* stimulus conditions *i* shown in **Figure 3** (Dataset 1). The third and fourth terms correspond to fitting the gain and phase of eye velocity (E) during the VOR before learning for each of the *m* frequencies *j* in Dataset 2 (**Figure 6A**). The fifth and sixth terms correspond to enforcing the gain and phase of Purkinje cell activity during the VOR at 0.5 Hz from Dataset 1. The final two terms enforce the steady state Purkinje cell (*P*_SS_) activity and eye velocity (*E*_SS_) during the VOR (extracted from the 500 ms step in head velocity in Dataset 1).

*C*_PostLearn_ was calculated for both increases and decreases in VOR gain and summed together with *C*_PreLearn_ to produce the total cost. The first two terms of *C*_PostLearn_ correspond to fitting the *changes* in gain and phase of eye velocity during the VOR over learning for each frequency in Dataset 2. The third and fourth terms correspond to enforcing the *changes* in the gain and phase of Purkinje cell activity during the VOR at 0.5 Hz (Dataset 1 and Datasets 3 and 4). The fifth term encourages the transient Purkinje cell responses to steps 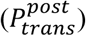 to mimic the transient responses after learning reported in Dataset 3 (Lisberger et al., 1994). The final two terms enforce the steady state Purkinje cell activity and eye velocity during the VOR after learning (Datasets 3 and 4).

Next, the visual filters at baseline (*k*_PR_ and *k*_PT_) were fine-tuned using the Nelder-Mead simplex method (MATLAB function *fminsearch*) to minimize the cost function:

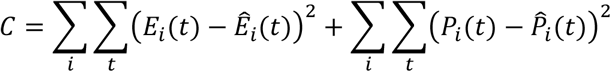

Here, *i* refers to the subset of stimulus conditions shown in **Figure 3** (Dataset 1) that include visual input (all conditions except VOR in the dark).

The models were fit to reproduce learned changes in Purkinje cell firing rate over learning only during the VOR, leaving changes in firing rate during cancellation as a prediction of the model. However, previous studies show that the amplitude of eye movements during VOR cancellation at low frequencies (0.5 Hz) is unchanged after learning (Guo & Raymond, 2010). To replicate this observation and ensure that the motor performance during VOR cancellation at low frequencies did not change after learning, we assumed that the visual prediction pathway *k*_PT_ contributed to maintenance of constant VOR cancellation performance after learning. This was enforced by adjusting *k*_PT_ after learning to minimize the cost function:

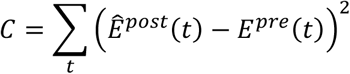

where *E* represents the eye velocity during VOR cancellation (x0 condition, see **Figure 3**) for 0.5 Hz sinusoidal stimuli in normal (pre) or high (post) gain monkeys. We emphasize that allowing different values for *k*_PT_ after learning does not imply that plasticity is required in the visual pathway. Instead, the changes in the visual prediction signal reflect the ability of primates to accurately anticipate the necessary visual tracking command to follow motion of a repeatable stimulus, after accounting for the gain of the VOR.

Simulations were run by calculating the convolutions in equations (1) and (2) numerically with 0.5 ms time steps. In **Figure 3C**, the root mean square error (in units of sp/s or °/s) was normalized by dividing the model error by the maximum stimulus speed (in units of °/s) during each behavioral condition and then averaging across conditions.

### 8.3 Schematic of idealized and “biological” linear temporal filters

The schematics illustrating idealized and “biological” linear temporal filters in **Figure 4C** are for pedagogical purposes, to aid in the interpretation of the model filters and step responses in **Figure 4B,C**. The idealized filters were convolved with a step function to produce the idealized step response. The biological filters for acceleration and velocity were constructed by convolving the idealized filters with an exponential filter, twice. The biological filter for position was constructed manually. All biological filters were then convolved with a step function to produce the “biological” step response.

### 8.4 Model analysis

To mathematically illustrate the degeneracy of the VOR circuit, we analyzed a slightly simplified version of the model with no explicit visual prediction pathway or explicit consideration of delays (which would provide a linear delay element), so that the filter components at all complex frequencies *s* obey the same equations in the Laplace domain:

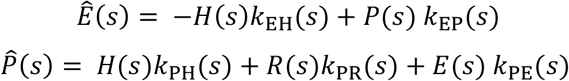

This allowed us to solve for the model parameters analytically in closed form. Further, the cost function of the model for a given dataset at steady state could be easily visualized as subsets of parameters were varied systematically (**Figure 5** and **Figure 8A**; steady state gain approximated by the 0.5 Hz sinusoidal conditions from Dataset 1, with the gain of efference copy positive feedback fixed at *g* = 0.2). In these plots, for ease of visualization, we display results for the steady state response (*s* = 0), for which the imaginary parts of the filters equal zero. Because the model is linear, all other stimulus conditions can be constructed as linear combinations of the VOR in the dark condition (vestibular input stimulus only) and the smooth pursuit condition (visual target stimulus only). This gives the following equations for VOR in the dark:

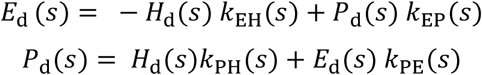

and for smooth pursuit:

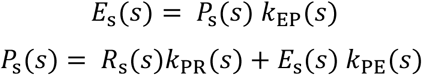

where *E*_d_ and *P*_d_ represent the complex gain and phase of the eye and Purkinje cell firing rate, respectively, during VOR in the dark, and *E*_s_ and *P*_s_ represent the same during smooth pursuit. Solving the model equations above shows that the brainstem parameters are fully constrained, and can be determined directly from the data as follows (equations below hold separately for every value of *s*):

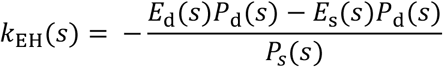

and

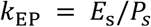

The parameters describing the inputs to the Purkinje cells (*k*_PH_, *k*_PE_, *k*_PR_) are degenerate, but related to each other by the following equations:

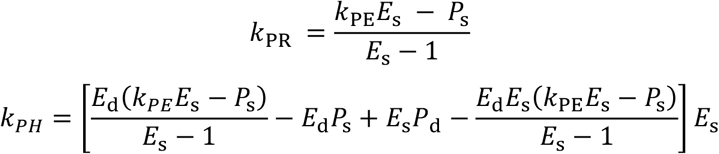

Thus, given the filter describing one pathway, the other two pathway’s filters can be determined. This observation motivated our strategy of fixing the efference copy filter pathway at different levels of feedback, and observing the requirements this imposes upon the other two pathways.

We also analyzed the model with a Purkinje cell “stimulation” condition conducted with the head stationary in the dark to mimic experiments in which Purkinje cells were electrically stimulated (Lisberger, 1994; **Figure 8C**). This provides one additional constraint, making the model nondegenerate when the stimulation condition is included. For this condition, the Purkinje cell equation was modified to include an additional input *S(s)* that was only delivered to Purkinje cells:

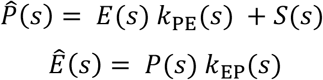

During this condition, the Purkinje cell response derived from these two equations is:

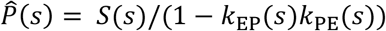

### 8.5 Simulation of paradoxical changes

To simulate the “paradoxical” changes in neural activity during VOR cancellation and smooth pursuit following learning (Miles and Lisberger, 1981; Lisberger, 1994; **Figure 7A,E**), we expanded our model to create a population of 20 Purkinje cells whose average activity matched the model fits in the remainder of the paper. For each “cell”, *k*_PH_ and *k*_PR_ were scaled by a random factor drawn from a normal distribution *N*(1, σ) with σ = 0.25 and 0.4, respectively, for the No Feedback model, and σ = 0.1 and 20 for the Strong Feedback model. These random scale factors were normalized to ensure that their average across the population for each model was exactly equal to one, which was necessary to ensure stability in the Strong Feedback model. We then examined the predicted Purkinje cell activity during smooth pursuit and during VOR cancellation before and after simulated learning. The amplitude of the Purkinje cell responses during VOR cancellation (sinusoidal stimulation at 0.5 Hz) was taken as the “Head Sensitivity” of the cell, mimicking the process used to estimate head sensitivity experimentally (Lisberger, 1994; Miles & Lisberger, 1981). Similarly, to simulate the process used to estimate “Eye Sensitivity” experimentally, the amplitude of the Purkinje cell responses to visual input alone was measured. “Head Sensitivity” was plotted against “Eye Sensitivity” to determine whether each model could replicate the previous finding of apparent changes in Purkinje cell “Head Sensitivity” (for a given “Eye Sensitivity”) with changes in VOR gain.

### 8.6 Transient stimulation

To simulate the effect of transient perturbation of Purkinje cells while the head was stationary in the dark (Lisberger, 1994; **Figure 8B**), we added a direct input signal of constant amplitude to the model Purkinje cell for 25 ms, and then allowed Purkinje cell activity and eye velocity to evolve according to the model dynamics (**Figure 8C**). Eye velocity outputs were scaled to be of equal maximum amplitude across efference copy feedback strengths.

## 9 Supplemental figures

**Figure 3–Figure Supplement 1:**
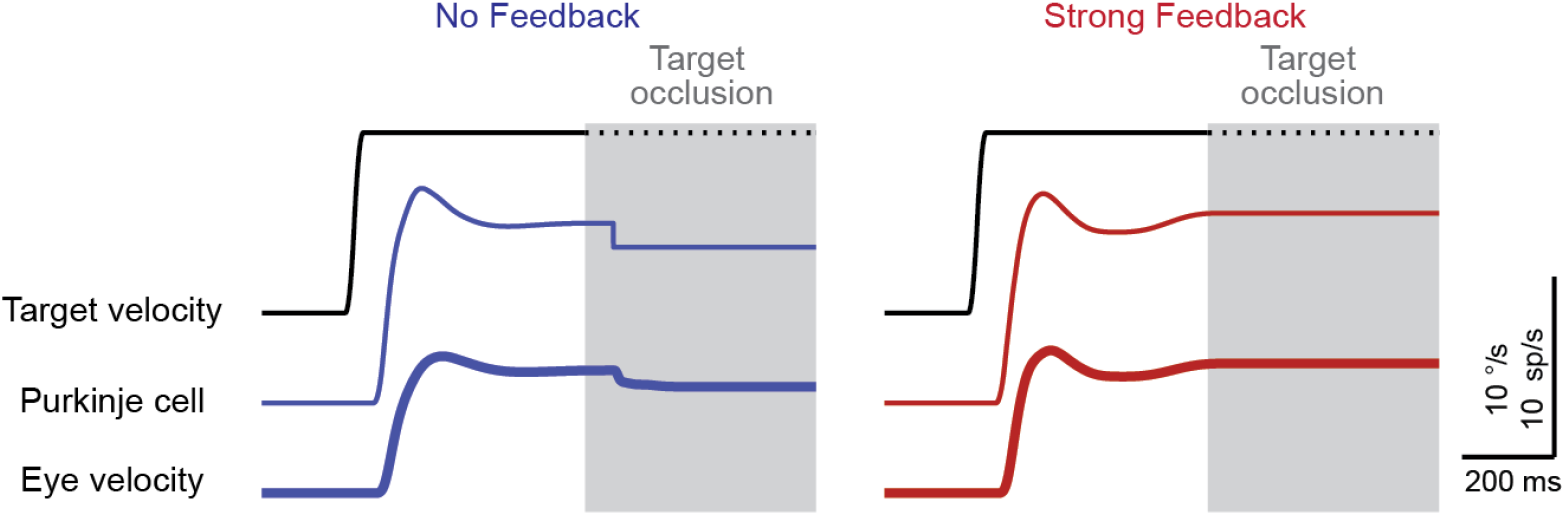
Model response to occlusion of a moving visual target. Target occlusion was simulated in each model by providing a step in target velocity as input, and once steady-state smooth pursuit was reached, removing all visual input. Efference copy feedback in the cerebellum has been proposed to be critical for maintaining ongoing smooth pursuit during steady-state tracking when retinal slip is minimal (Lisberger 1994), when the target is artificially stabilized on the retina (Stone & Lisberger, 1990), or when the eyes continue tracking a moving target that transiently disappears from view behind an occluder (Becker & Fuchs, 1985; Leung & Kettner, 1997; Stone & Lisberger, 1990). During transient target disappearance, humans and nonhuman primates continue tracking the invisible target with eye velocity between 40-100% of baseline (Becker & Fuchs, 1985; Bennett & Barnes, 2003; Churchland et al., 2003; Madelain & Krauzlis, 2003). Both the No Feedback and Strong Feedback models displayed maintained pursuit velocity during simulated target occlusion, with eye velocity perfectly maintained in the Strong Feedback model and partially maintained in the No Feedback model.

**Figure 3–Figure Supplement 2:**
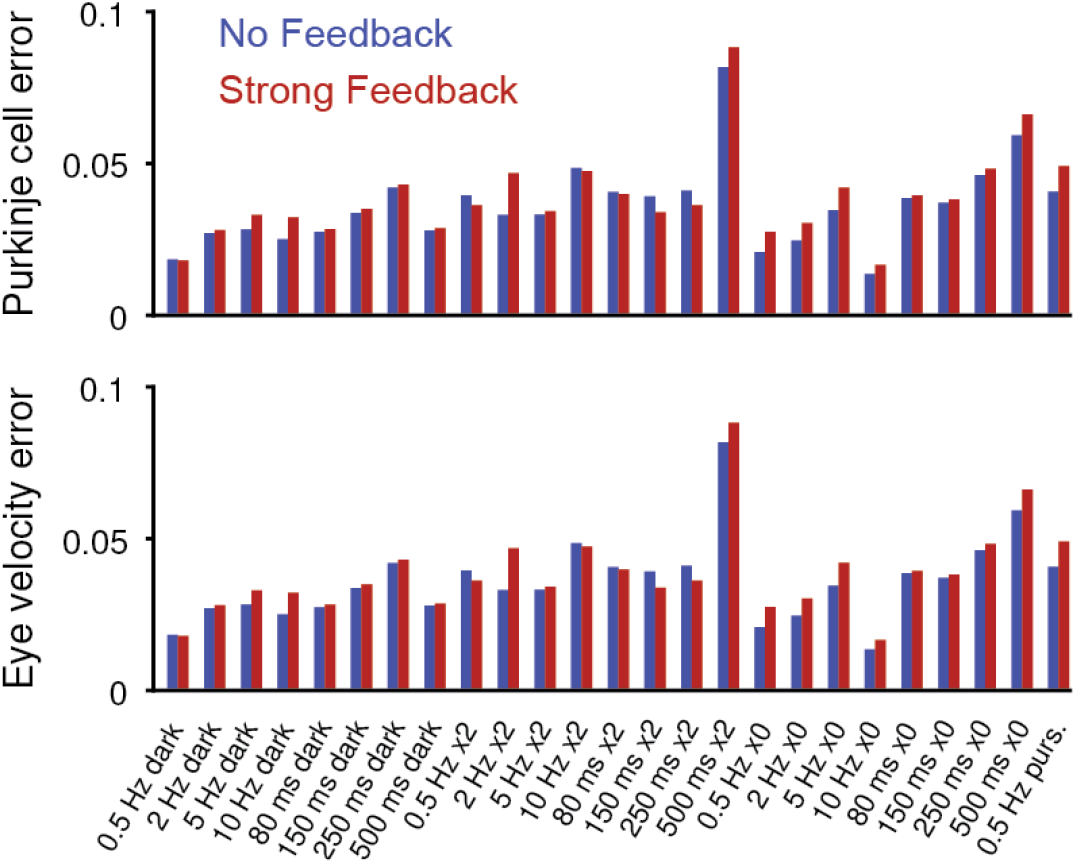
Model error for individual behavioral conditions. Purkinje cell error (*top*) and eye velocity error (*bottom*), for the No Feedback (*blue*) and Strong Feedback (*red*) models, for each stimulus condition shown in **Figure 3**.

**Figure 6–Figure Supplement 1:**
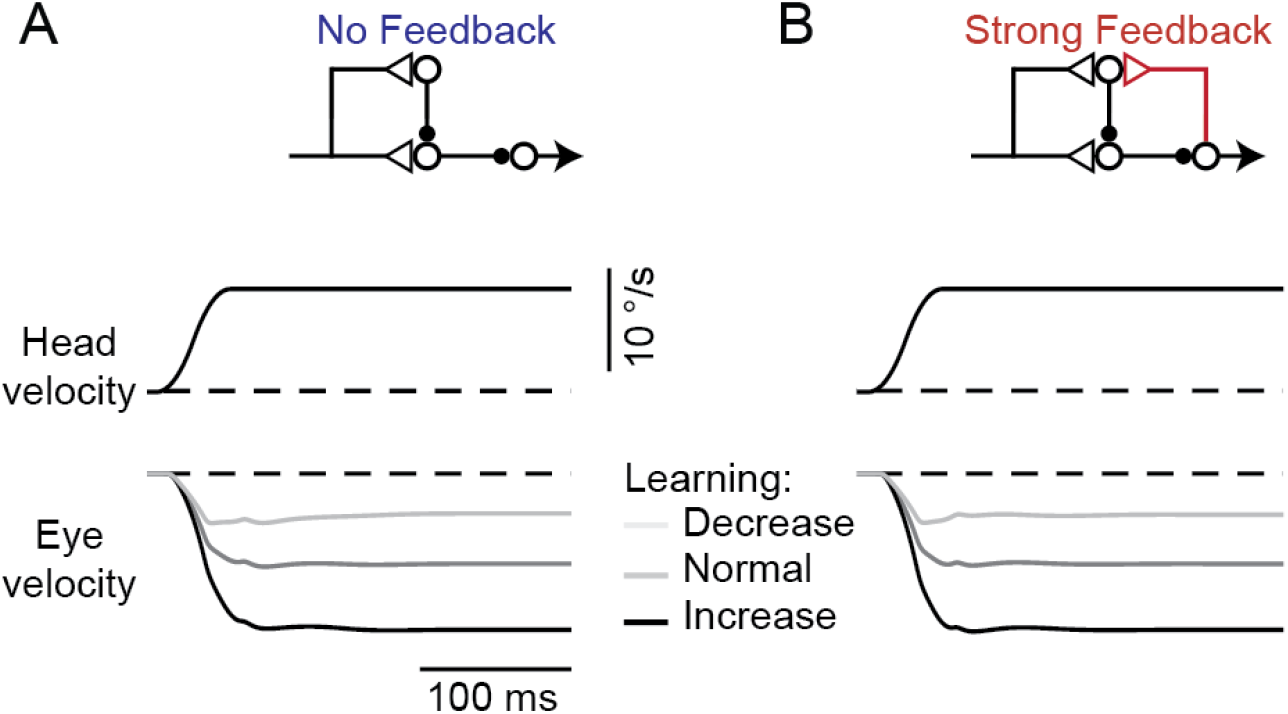
Model response to steps in head velocity after VOR learning. **(A,B)** Model eye velocity responses to steps in vestibular input (VOR in the dark), before and after learning in the No Feedback model (A) and the Strong Feedback model (B), both before learning (“Normal”) and after learned increases or decreases in VOR gain.

**Figure 6–Figure Supplement 2:**
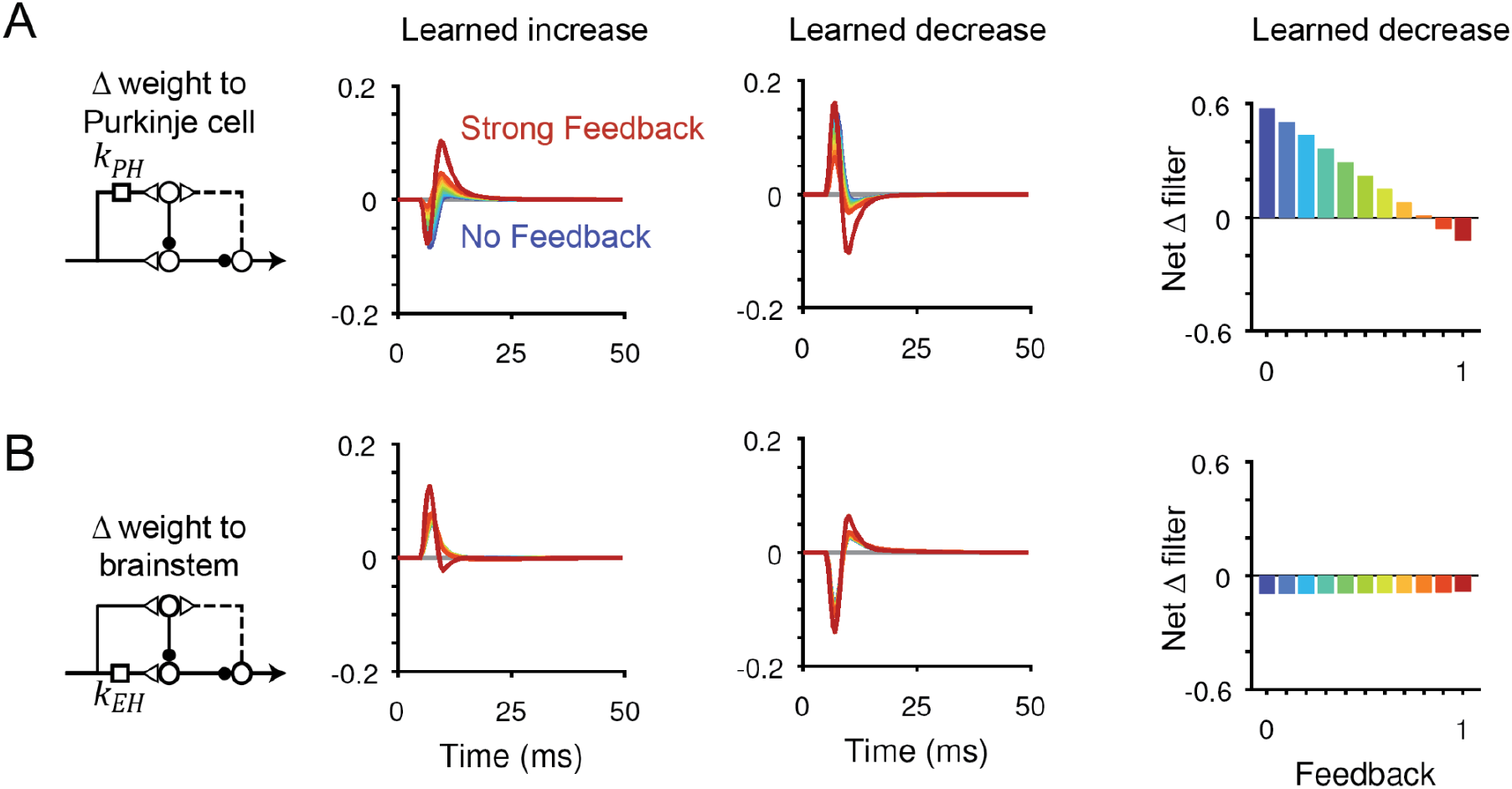
Changes in filter strength after VOR learning for all models. **(A)** Dynamic changes in the filter *k*_PH_ after learned increases (*left*) or decreases (*center*) in VOR gain. *Right*, net change in filter strength after learned decreases in VOR gain. **(B)** Same as (A) but for the filter *k*_EH_.

**Figure 6–Figure Supplement 3:**
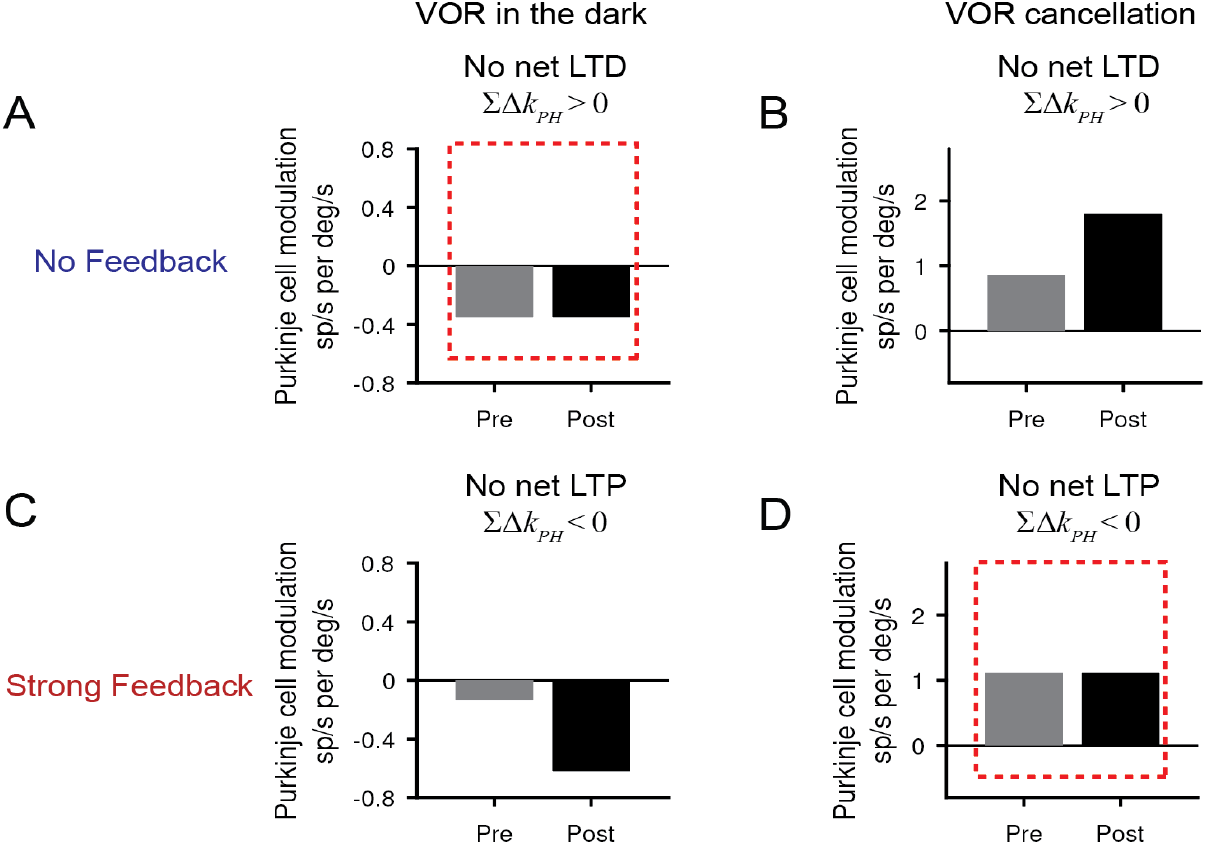
Changes in neural activity after learning when the direction of plasticity in the cerebellar cortex is restricted. **(A)** Purkinje cell modulation during the VOR in the dark in the No Feedback model when LTD was blocked. To computationally block LTD, the net change in filter *k*_PH_ was restricted to be positive (“No net LTD”). Without LTD, Purkinje cell activity does not decrease after learned increases in VOR gain (compare to **Figure 6–Figure Supplement 4A**). **(B)** Purkinje cell modulation during VOR cancellation in the No Feedback model when LTD was blocked. The increase in Purkinje cell modulation during VOR cancellation does not require LTD. **(C)** Same as (A) but for the Strong Feedback model with LTP blocked. For this model, the change in Purkinje cell modulation during the VOR does not require LTP. **(D)** Same as (B) but for the Strong Feedback model. Without LTP, Purkinje cell activity does not increase during VOR cancellation after learning.

**Figure 6–Figure Supplement 4:**
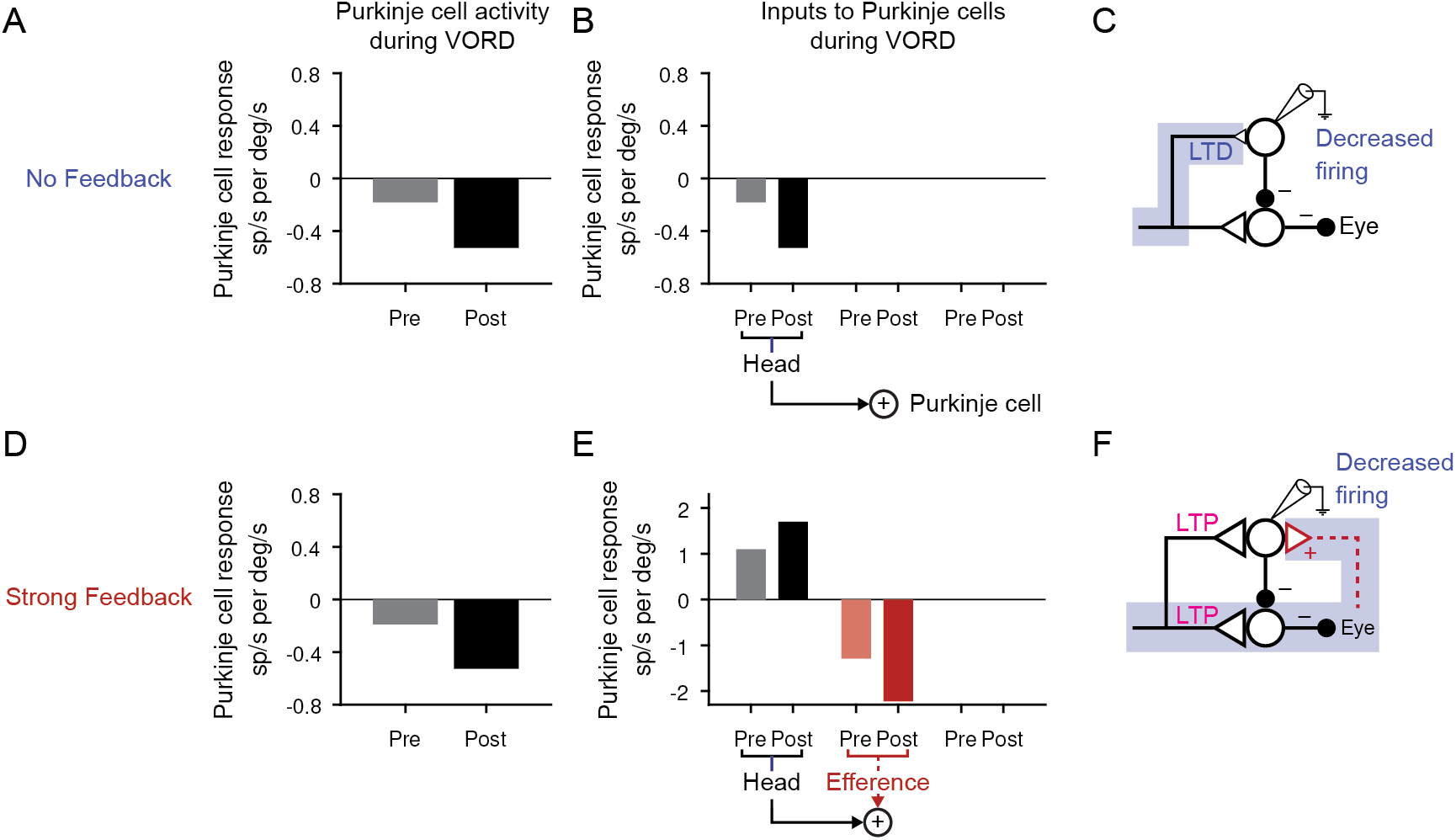
Changes in Purkinje cell activity during the VOR in the dark. **(A)** Purkinje cell firing rate response during the VOR in the dark, before (“Pre”) and after simulated learned increases (“Post”) in the gain of the VOR for the No Feedback model, during 0.5 Hz vestibular input. Purkinje cell response was plotted as negative if the neural activity was >90° out of phase with the ipsiversive stimulus. Compare to results in (Watanabe, 1985) (their Figure 6); (Lisberger et al., 1994) (their Figure 10); (Hirata & Highstein, 2001) (their Figure 3), and (Blazquez et al., 2003) (their Figure 2C). **(B)** Source of Purkinje cell modulation during the VOR for the No Feedback model. Each bar represents the amplitude of a 0.5 Hz sinusoidal signal as it reaches the Purkinje cell after passing through the relevant filter in spikes/s, normalized to the amplitude of the head velocity stimulus in °/s. The only input to Purkinje cells in this condition is the head velocity pathway. **(C)** Schematic highlighting the path responsible for the observed change in Purkinje cell firing. In the No Feedback model, the observed change in neural activity is due to plasticity in feedforward pathways. **(D)** Same as (A) for the Strong Feedback model. **(E)** Same as (B) for the Strong Feedback model. The potentiation in the head velocity input pathway is counteracted by the large decrease in the efference copy pathway input to the Purkinje cells. **(F)** In the Strong Feedback model, the observed decrease in Purkinje cell activity is driven by plasticity in the brainstem, fed back to Purkinje cells through the efference copy pathway.

**Figure 7–Figure Supplement 1:**
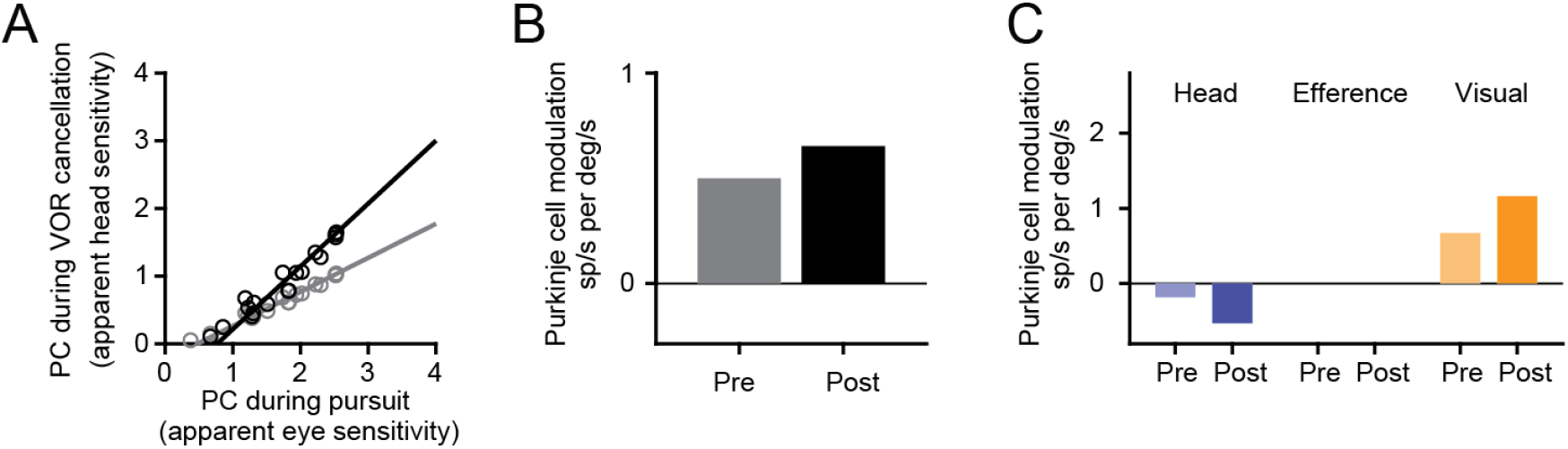
Explanation for changes in Purkinje cell activity during VOR cancellation in a model without a visual prediction pathway. Paradoxical changes during VOR cancellation are still reproduced by the No Feedback model without a visual prediction pathway. See legend for **Figure 7A,B,C**.

**Figure 8–Figure Supplement 1:**
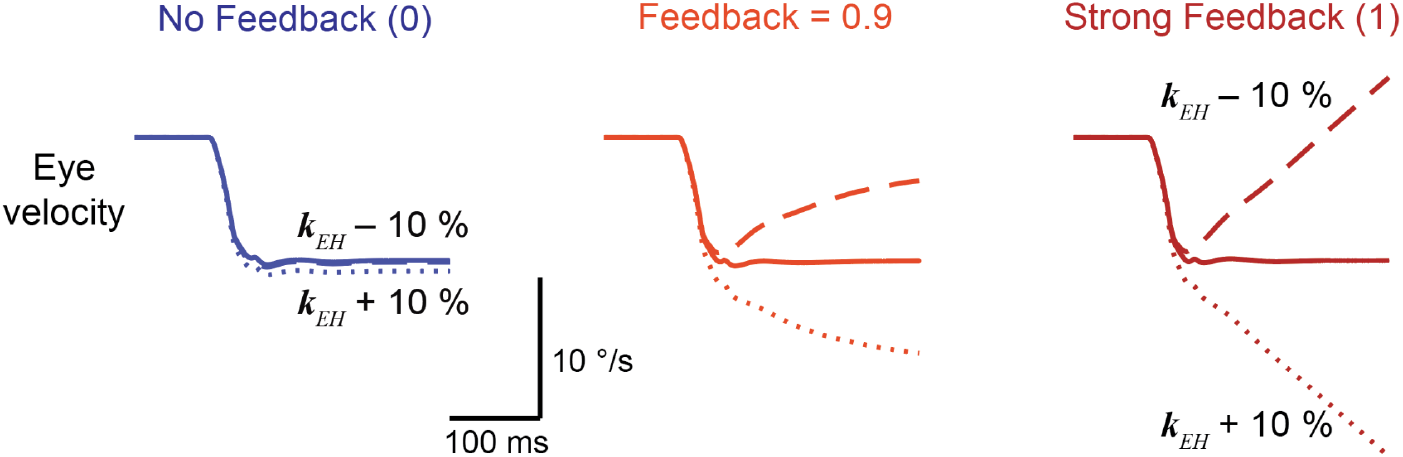
Sensitivity of models to parameter changes. Model output showing Purkinje cell activity (*top*) and eye velocity (*bottom*) in response to an ipsiversive step in head velocity, for No Feedback, Feedback = 0.9, and Strong Feedback models. Results are shown for the original models before learning (*solid*), and after increasing (*short dash*) or decreasing (*long dash*) the amplitude of the filter carrying vestibular input to the brainstem (*k*_EH_) by 10%.

